# Population structure and genomic evidence for local adaptation to freshwater and marine environments in anadromous Arctic Char (*Salvelinus alpinus*) throughout Nunavik, Québec, Canada

**DOI:** 10.1101/2020.04.29.066449

**Authors:** Xavier Dallaire, Éric Normandeau, Julien Mainguy, Jean-Éric Tremblay, Louis Bernatchez, Jean-Sébastien Moore

## Abstract

Distinguishing neutral and adaptive genetic variation is one of the main challenges in investigating processes shaping population structure in the wild, and landscape genomics can help identify signatures of adaptation to contrasting environments. Arctic Char (*Salvelinus alpinus*) is an anadromous salmonid and the most harvested fish species by Inuit people, particularly so in Nunavik (Canada), one of the most recently deglaciated region in the world. Unlike most other anadromous salmonids, Arctic Char occupy coastal habitats near their overwintering rivers during their marine phase. The main objective of this study was to document the putative neutral and adaptive genomic variation of anadromous Arctic Char populations in Nunavik (Québec, Canada) and bordering regions. A second objective was to interpret our results in the context of fisheries management in Nunavik. We used genotyping-by-sequencing (GBS) to genotype 18,112 filtered single nucleotide polymorphisms (SNPs) for 650 individuals sampled in 23 locations (average sample size per location = 28) along >2,000 km of coastline. Our results reveal a hierarchical genetic structure, whereby neighboring hydrographic systems harbour distinct populations grouping within major oceanographic basins, namely the Hudson Bay, Hudson Strait, Ungava Bay and Labrador Sea. We found genetic diversity and differentiation to be consistent with both the expected post-glacial recolonization history and patterns of isolation-by-distance reflecting contemporary gene flow. Furthermore, using three gene-environment association (GEA) methods we found genomic evidence for local adaptation to freshwater and marine environmental components, especially in relation to sea-surface and air temperatures during summer, as well as salinity. Our results support fisheries management at a regional level, and other implications on hatchery projects and adaptation to climate change are discussed.

## 1. Introduction

Intraspecific diversity is an important part of biodiversity, especially at higher latitudes where species richness is relatively low (Pamilo & Savolainen, 1999). In northern regions, the periodic range contractions and expansions brought about by glacial cycles have shaped this diversity through bottlenecks and genetic drift (Hewitt, 2000). Local adaptation can also arise when species experience different environmental conditions over their geographic ranges (Kawecki & Ebert, 2004; Williams, 1966). Local adaptation has been studied extensively via reciprocal transplant and common-garden field experiments, but these approaches do not provide information on the molecular basis of adaptation (Tiffin & Ross-Ibarra, 2014). New genomic methods are now commonly used to advance our understanding of local adaptation (Grummer et al., 2019; Luikart et al., 2018). Such adaptive genomic variation, as well as contemporary population genetic structure, are of great interest for both conservation and management to ensure actions target biologically significant units (Bernatchez et al., 2017; Funk, McKay, Hohenlohe & Allendorf, 2012).

Salmonids are a diverse family of fishes with high economic and cultural importance. Many populations have an anadromous life cycle, whereby individuals are born and reproduce in freshwater and migrate to the sea to feed and grow. Anadromous salmonids are philopatric, i.e., returning to their natal habitat for spawning (Quinn, 1993) and this behaviour is known to reduce gene flow amongst populations, thus promoting genetic differentiation and local adaptation at fine spatial scales (Fraser, Weir, Bernatchez, Hansen & Taylor, 2011). Recent studies have identified genomic regions associated with many environmental parameters, including air temperature (Bourret, Dionne, Kent, Lien & Bernatchez, 2013; Hand et al., 2016; Perrier, Ferchaud, Sirois, Thibault & Bernatchez, 2017; Sylvester et al., 2018), precipitation (Hecht, Matala, Hess & Narum, 2015; Micheletti, Matala, Matala & Narum, 2018), geology (Bourret et al., 2013), upstream catchment area (Pritchard et al., 2018) and migration distances (Hecht et al., 2015; Micheletti et al., 2018) in many species of salmon and trout. Little is known, however, about the genomic basis of marine adaptations for these species even though they spend a significant proportion of their lives at sea. The capacity to identify divergent selective pressures on salmonid populations at sea is mostly limited by our understanding of their geographically extensive and highly variable marine distribution (Hecht et al., 2015; Quinn, 2005).

The Arctic Char (*Salvelinus alpinus*) is a salmonid fish with a circumpolar distribution and is known for its great diversity in life-history characteristics (Klemetsen, 2010). Anadromous individuals spend 3 to 9 years in cold oligotrophic freshwater at birth (Johnson, 1980), then complete annual migrations between marine habitats for summer foraging and lakes for overwintering. Although straying (i.e., an upstream migration in a non-natal river system) can occur, several studies have shown that Arctic Char maintained philopatric behaviour during reproductive years, limiting effective dispersal (Gyselman, 1994; Moore, Harris, Tallman & Taylor, 2013; Moore et al., 2017). Moore et al. (2013) also argued that gene flow could be sufficiently low to allow for local adaptation among populations of eastern Baffin Island in the Canadian Arctic, while Moore et al. (2017) provided some genomic evidence for local adaptation to natal rivers at a fine spatial scale. In marine environments, Arctic Char tend to stay near the surface (< 3m), with occasional dives up to 50 m (Harris et al., 2020; Spares, Stokesbury, O’Dor & Dick, 2012) and preferably use nearshore habitats within approximatively 100 km from their natal river’s mouth (Dempson & Kristofferson, 1987; Layton et al., 2020; Moore et al., 2016). Thus, one could expect this behaviour to lead to Arctic Char populations experiencing more contrasted marine conditions than other anadromous species.

During the last glacial maximum (LGM), around 18,000 years ago, most of actual Canada was covered by glaciers (Dyke, 2004), contracting species’ ranges to ice-free glacial refugia. Potentially strong demographic bottlenecks during glaciations have greatly reduced intraspecific genetic diversity and lead to divergent glacial lineages that survived in different refugia (Hewitt, 2000). Secondary contact between lineages have been shown to have commonly occurred among temperate freshwater fishes (e.g., Atlantic Salmon *Salmo salar*: Bradbury et al., 2015; Lake Cisco *Coregonus artedi*: Turgeon & Bernatchez, 2001; Lake Whitefish *Coregonus clupeaformis*: Bernatchez & Dodson, 1990). In North America, Arctic Char comprises four known mitochondrial DNA (mtDNA) lineages associated with distinct glacial refugia (Brunner, Douglas, Osinov, Wilson & Bernatchez, 2001; Moore, Bajno, Reist & Taylor, 2015). The Arctic lineage is predominant in the south-eastern Arctic (Brunner et al., 2001, Moore et al., 2015), but populations across northern Labrador are admixed with the Atlantic lineage (Salisbury, McCracken, Keefe, Perry & Ruzzante, 2019).

Nunavik, situated in northern Québec (Canada), is one of the last regions in North America to have deglaciated following the LGM (Dyke, 2004). It is bordered by the Hudson Bay, Hudson Strait, and Ungava Bay (Figure 1). These three marine regions are contrasted in their surface temperature, salinity, productivity, and tidal regimes (Prisenberg, 1984; Savard et al., 2014). In remote coastal communities of Nunavik, Arctic Char subsistence fisheries are key to food security (Laflamme, 2014) and rapid demographic growth has increased harvesting pressures on wild fish populations (Martin, 2011). The current study applies population genomic methods to investigate the relative role of neutral vs. adaptive processes in shaping contemporary population structure of anadromous Arctic Char over 2,000 km of coastline in Nunavik. This will provide a genomic context of relevance for sustainable fisheries management.

**Figure 1:**
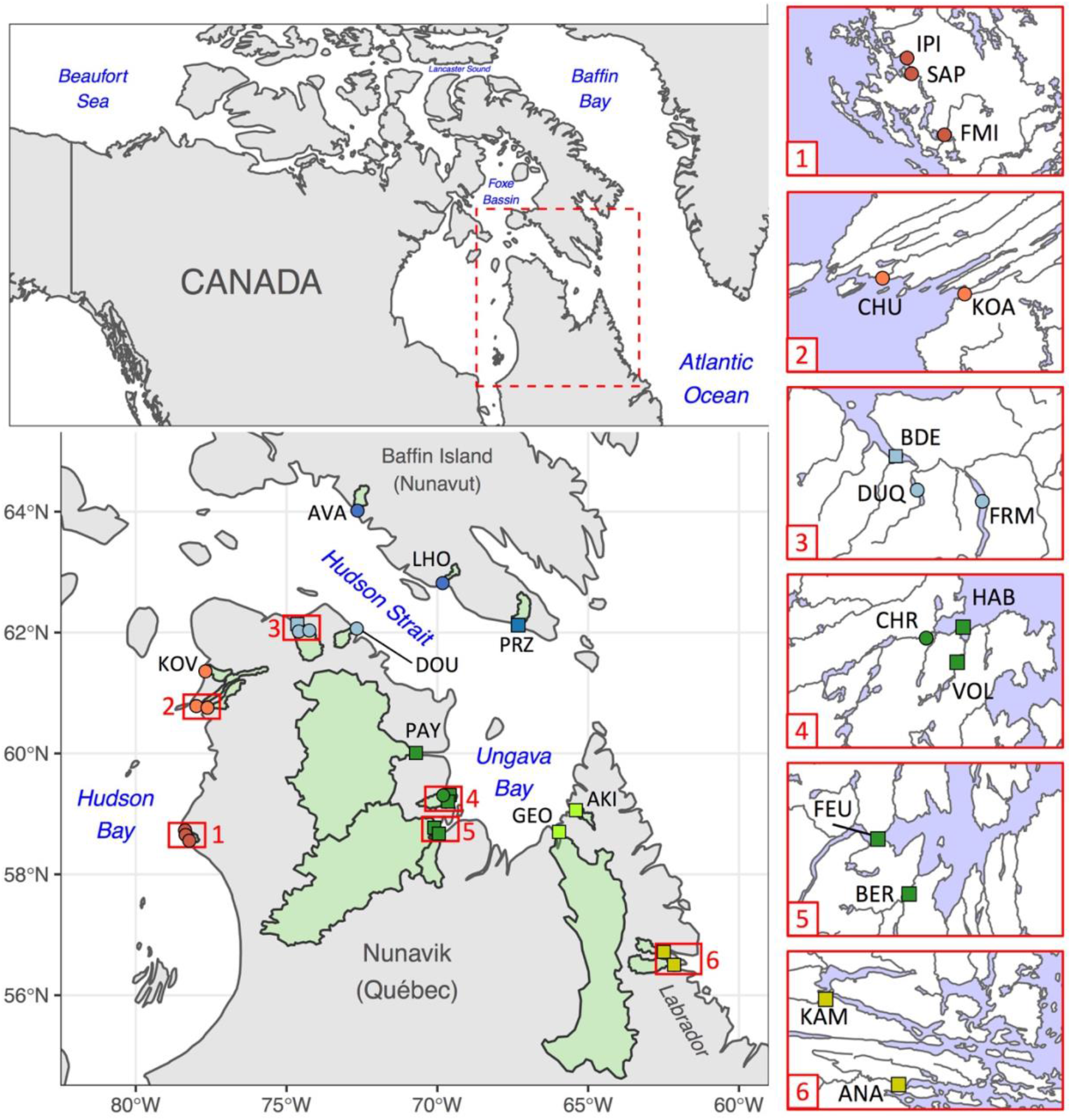
Sampling locations in Nunavik (Québec) and bordering regions. Red extents numbered 1 to 6 are magnified to show neighboring sampling locations. Samples were either used for Genotyping-by-Sequencing (GBS) only (round) or for GBS and mitochondrial DNA sequencing (square). Catchment area for each site is displayed in green.

The entire area surveyed in this study was covered by ice from the LGM up to approximately 9,000 – 11,000 years ago (Dyke, 2004). As Arctic Char are thought to have rapidly recolonized habitats following the retreat of the glaciers (Hammar, 1987; Power, Pope & Goad, 1973) and have relatively long generation time (Gulseth & Nilssen, 2001), we expected the contemporary genetic structure of Arctic Char populations in Nunavik to be heavily influenced by their recent post-glacial recolonization history. Specifically, we first sequenced the mitochondrial control region to identify glacial lineages present in Nunavik to place the present study in the context of previous work in the region that relied heavily on mtDNA. We then used genotyping-by-sequencing (GBS) to document Arctic Char neutral genetic structure in relation to its post-glacial history by i) delimiting putative populations via a clustering analysis, ii) comparing levels of genetic diversity between geographic regions, and iii) investigating patterns of isolation-by-distance and the importance of potential barriers to migration. Finally, since we expected broad-scale variation in both freshwater and marine environments to be the source of divergent selective pressures, we attempt to detect evidence of local adaptation in Arctic Char around Nunavik and bordering regions. We extracted various environmental variables representing the different habitats occupied by Arctic Char and combined multiple gene-environment association methods to identify candidate SNPs associated with environmental factors.

## 2. Methods

### 2.1 Sampling and DNA extraction

Arctic Chars were sampled in 23 water bodies across Nunavik, southern Baffin Island, and Labrador (Figure 1). In Nunavik and Baffin Island, adult fish were harvested during their upstream migration using gillnets or counting weirs. In most localities throughout Nunavik, sampling locations were selected in concert with local and regional Inuit wildlife managers, and sampling was done with the assistance of local Inuit guides, to prioritize fish populations with an importance for traditional fishing. In two locations (Deception Bay and Hopes Advance Bay), samples were taken in the estuary as well as in two tributary rivers. Juvenile fish were captured by electrofishing in two rivers near Nain, Labrador. Adipose fin clips were collected from each fish and preserved in ethanol 95%. DNA was extracted from fin clips using a modified version of Aljanabi & Martinez (1997). Agarose gel electrophoresis was used to assess DNA quality and the quantity and quality of DNA was evaluated by NanoDrop spectrophotometer (Thermo Scientific).

### 2.2 Mitochondrial DNA analysis

Based on the analysis of mtDNA, Salisbury et al. (2019) showed an overlap of the Arctic and Atlantic lineages in Northern Labrador (55 – 59° latitude), and an absence of correlation between latitude and lineage prevalence which suggests a complete introgression of fish from Arctic and Atlantic origins in the region. To assess whether the two glacial lineages also hybridized in Nunavik, which could influence interpretations of nuclear diversity results, we sequenced mtDNA for a subset of individuals from sampling sites in the eastern part of our study area (n = 10 to 34 per site). The entire mitochondrial control region was amplified with primers Tpro2 (Brunner et al., 2001) and SalpcrR (Power et al., 2009) and we sequenced 499 base pairs of the control region left domain region according to methods outlined in Power et al. (2009). Control region sequences were aligned on the reference haplotype set described in Salisbury et al. (2019) using the geneious (6.1.8; www.geneious.com) alignment tool with parameters set to default and the closest matching haplotype was reported.

### 2.3 GBS library preparation, sequencing and SNP calling

Ten (10) μl of DNA samples were normalized to a concentration of 10 ng/μl using Quant-iT Picogreen dsDNA Assay Kit (Invitrogen) for precision quantification. GBS libraries were prepared with a modified version of the Abed et al. (2019) two-enzyme GBS protocol, using *Pst*I and *Msp*I restriction enzymes. Samples were randomly assigned to libraries to limit batch effects. Sequencing was done on Ion Torrent p1v3 chips with a median target of 80 million single-end reads per chip. Each library was sequenced on 3 separate chips, and the volume of DNA from each sample was adjusted after the first chip to reduce the unbalanced representation of individuals in sequences.

We processed the data and filtered the SNP dataset using a RADseq workflow (https://github.com/enormandeau/stacks_workflow) built around STACKS 2.5 (Rochette, Rivera-Colón & Catchen, 2019). Briefly, the sequences were trimmed at 80 bp and aligned on a *Salvelinus* sp. reference genome (ASM291031v2; NCBI RefSeq: GCF_002910315.2; Christensen et al., 2018) using bwa mem (-k 19 -c 500 -O 0,0 -E 2,2 -T 0) in BWA-0.717 (r1188; Li & Durbin, 2009) and samtools view (-Sb -q 1 -F 4 -F 256 -F 2048) in samtools 1.8 (Li et al., 2009). SNPs were called on polymorphic genotypes with at least 4X coverage per sample (-m), present in at least 60% of samples of each sampling sites (-r) and with the minor allele present in a minimum of 3 samples. This minor allele sample (MAS) filter is akin to minor allele frequency (MAF), but unlike MAF it is not artificially boosted by frequent RADseq genotyping errors (Linck & Battey, 2019). Samples with more than 20% of missing data or with heterozygosity (F_IS_) under -0.2 were removed. For SNP calling and quality control, we combined our samples with 119 other samples from three sampling sites that weren’t included in our final dataset, as they either had too few samples, had no access to sea, or targeted a population reintroduced from a hatchery brood.

Salmonid fishes have a common ancestor that experienced a whole-genome duplication approximately 60 MYA (Crête-Lafrenière, Weir & Bernatchez, 2012), and many genetic markers identified in our analyses are expected to be situated on paralogous loci of similar sequences. While these loci may be important for adaptation (Kondrashov, 2012), they were removed due to the fact that these markers do not behave like bi-allelic SNPs and because genotyping is difficult without very high coverage (>100 reads; Dufresne, Stift, Vergilino & Mable, 2014). SNPs on duplicated loci were categorized and filtered using an adapted *HDplot* procedure (McKinney, Waples, Seeb & Seeb, 2017), which identifies paralogs by visually comparing the allelic ratio, the proportion of heterozygotes and homozygotes of the rare allele, and the F_IS_ value for each SNP.

We also filtered the markers to avoid physically linked SNPs while keeping a maximum of information: for each pair of SNPs within 50,000 bp, we assessed linkage considering samples without missing data where at least one of the two genotypes contains the rare variant. If the two markers had identical genotypes in more than 50% of these samples, the pair were considered linked and only the first SNP was kept. To identify markers linked to sex, we performed a redundancy analysis (RDA) using individual genotypes from samples for which information on sex was available in Hudson Bay (n = 69) and Ungava Bay (n = 85). We classified SNPs as showing statistically significant association with sex when they loaded with more than 2.5 standard deviation from the mean (p = 0.012) and removed those markers from the dataset. Finally, we measured pairwise relatedness (Yang et al., 2010) in vcftools v0.1.13 (Danecek et al., 2011) to make sure that our samples did not include close siblings (relatedness > 0.5).

### 2.4 Identification of putative neutral markers

Markers potentially under selection were identified using two methods: pcadapt (Luu, Bazin & Blum, 2017) and BayeScan (Foll & Gaggiotti, 2008). SNPs identified as outliers by at least one method were removed to produce a neutral dataset. The R package pcadapt identifies outlier SNPs in relation to population structure using principal components analyses (PCA). The first 11 PCs were used, based on visual evaluation of PCA scores and scree plots, as recommended by pcadapt authors, and SNPs with minor allele frequencies under 0.05 were excluded from the analysis. We then used BayeScan with prior odds of 1,000 and other parameters set to default. For both tests, SNPs with a false detection rate (q-value) under 0.05 were considered putatively under selection.

### 2.5 Basic statistics and population structure

Population genetic statistics were computed from the neutral dataset. Observed and expected heterozygosity were calculated by population using GenoDive v3.0 (Meirmans & Van Tienderen, 2004). We used vcftools v0.1.13 to measure the proportion of heterozygous SNPs for each individual, and we calculated the number of polymorphic SNPs in each population. Effective population sizes (N_e_) and 95% confidence intervals were estimated with Neestimator v2.01 (Do et al., 2014) using the linkage disequilibrium method on markers with minor allele frequencies over 0.05.

We used a principal coordinate analysis (PCoA) in the R package *dartR* (Gruber, Unmack, Berry & Georges, 2018) to document population structure using the neutral data set. We also estimated ancestry with the maximum likelihood approach implemented in ADMIXTURE (Alexander, Novembre & Lange, 2009) with the number of genetic clusters (K) ranging from 1 to 20. We considered the value of K yielding the lowest cross-validation error to be the number of genetic groups best supported by ADMIXTURE. Based on this K value, we identified contiguous sampling sites where most individuals shared a common cluster membership and repeated the ADMIXTURE analysis within those sites with K ranging from 1 to 6. Considering that adult Arctic Char is expected to stray to nearby rivers, we regrouped sampling sites in connecting rivers and estuaries for the following analyses on genetic diversity and isolation-by-distance, with the exception of cases where ADMIXTURE showed strong population structure.

Spatial patterns of genetic diversity were compared between a priori regions, using the individual proportion of heterozygous SNPs. A nested ANOVA was performed with the regions as groups and the sampling sites as subgroups. To account for our unbalanced sampling design, we did a comparison of estimated marginal means (or least-square means) with the Satterthwaite approximation of degrees of freedom (Satterthwaite, 1946) on a linear mixed-effect model with a random effect of the sampling site, in the R packages *lme4* (Bates, Mächler, Bolker & Walker, 2015) and *emmeans* (Lenth, 2019).

### Landscape genomics

Pairwise population F_ST_ were calculated with the R package *StAMPP* (Pembleton et al., 2013), and 1,000 bootstraps were performed to estimate their significance value as the proportion of bootstrapped F_ST_ values under zero. F_ST_ calculations were performed on both the neutral and putatively adaptive datasets. The geographic marine distance separating sampling sites was measured between the coordinates at the mouth of sampled rivers by a least-cost path in the R package *marmap* (Simon-Bouhet, 2013), using NOAA bathymetric data at a 4-minutes resolution to discriminate land and sea.

We tested for the presence of isolation-by-distance (IBD) with a linear mixed-effect model. Linearized F_ST_ (F_ST_/(1-F_ST_)), based on the neutral dataset, was used as the dependent variable and marine distance as a fixed effect. Patterns of IBD between pairs of sampling sites on the same coast versus pairs on either side of Hudson Strait were compared by the inclusion of a categorical fixed effect. To characterize the impact of Hudson Strait as a barrier to gene flow at different spatial scales, analyses were performed on all non-estuarine sites, then only on non-estuarine sites within 250 km of the Hudson Strait. To account for the non-independence of pairwise distances, the model was run with a maximum likelihood population effect parameterization (MLPE) (Clarke, Rothery & Raybould, 2002) with and without restricted maximum likelihood (REML), using the MLPE.lmm function in the R package *ResistanceGA* (Peterman, 2018). Models without REML were compared with conditional Akaike Information Criterion (cAIC; Vaida & Blanchard, 2005) in the R package *cAIC4* (Säefken, Ruegamer, Baumann & Kruse, 2019), and with marginal R^2^ (Nakagawa & Schielzeth, 2013) in the R package *MuMIn* (Barton, 2019) for models with REML.

### 2.7 Gene-environment association

We used the ArcGIS software v10.4 (ESRI, 2011) to extract environmental data from BIO-Oracle v2.0 (Assis et al., 2018), Marspec (Sbrocco & Barber, 2013), and WorldClim v2.0 (Fick & Hijmans, 2017) (Table 1, see Table S5 for values at sites). Tide data were obtained from FES2014, produced by Noveltis, Legos and CLS and distributed by Aviso+, with support from Cnes (https://www.aviso.altimetry.fr/). Marine variables represent sea-surface values for factors of potential biological importance for Arctic Char and were aggregated in a 20 km radius around each river mouth and within 5 km from the coast, as to best represent the local coastal environment based on existing knowledge of Arctic Char marine habitat use from other geographical regions (Spares, Stokesbury, Dadswell, O’Dor & Dick, 2015; Moore et al., 2016). Freshwater variables comprise the area of the watershed upstream of the sampling site, as well as air temperature and precipitation statistics on these areas. Air temperature is commonly used as a proxy for freshwater temperature in remote areas, as supported by studies linking the growth rate of Lake Trout to air temperature (Black, von Biela, Zimmerman & Brown, 2013; Torvinen, 2017). To deal with co-linearity, a PCA was performed on the scaled environmental factors, and the PCs with eigenvalues over 1 were used as explanatory variables in gene-environment association (GEA) analyses.

**Table 1.** Value range and source of environmental factors considered in gene-environment associations. PCA axis most associated to variables are indicated with sign (positive or negative) of the correlation in parenthesis.

| Variable | Description | Value range | Unit | Temporal range | Database | Source | PCA axis |
| --- | --- | --- | --- | --- | --- | --- | --- |
| *Marine variables* |  |  |  |  |  |  |  |
| Productivity | Annual mean primary productivity (carbon) | 1.2 – 12.0 | g.m<sup>-3</sup>.day<sup>-1</sup> | 2000 – 2014 | Bio-ORACLE | PISCES | PC1 (+) |
| O2 | Annual mean dissolved molecular oxygen | 327 – 364 | mmol.m<sup>-3</sup> | 2000 – 2014 | Bio-ORACLE | PISCES | PC1 (-) |
| SST_summer | Mean sea-surface temperature (July to September) | 0.5 – 6.8 | °C | 2002 – 2010 | Marspec | WOA09 | PC2 (+) |
| Turbidity | Annual mean diffuse attenuation | 0.8 – 4.8 | m<sup>-1</sup> | 2002 – 2009 | Bio-ORACLE | Aqua-MODIS | PC2 (+) |
| Salinity | Annual mean sea-surface salinity | 21.0 – 31.8 | PSS | 2000 – 2014 | Bio-ORACLE | ARMOR | PC2 (-) |
| Tide_M2 | Amplitude of M2 tidal constituent | 17.5 – 372.6 | --- | --- | FES2014 | AVISO+ | PC3 (+) |
| *Freshwater variables* |  |  |  |  |  |  |  |
| P_winter | Mean precipitation of the coldest quarter | 40.2 – 171.5 | mm | 1970 – 2000 | WorldClim | GHCN | PC1 (+) |
| P_summer | Mean precipitation of the warmest quarter | 114.7 – 250.1 | mm | 1970 – 2000 | WorldClim | GHCN | PC1 (+) |
| T_winter | Mean air temperature of the coldest quarter | -25.0 – -19.7 | °C | 1970 – 2000 | WorldClim | MODIS | PC1 (+) |
| T_min | Minimum air temperature of the coldest month | -29.0 – -26.3 | °C | 1970 – 2000 | WorldClim | MODIS | PC1 (+) |
| T_summer | Mean air temperature of the warmest quarter | 4.9 – 10.7 | °C | 1970 – 2000 | WorldClim | MODIS | PC2 (+) |
| T_max | Maximum air temperature of the warmest month | 9.3 – 16.4 | °C | 1970 – 2000 | WorldClim | MODIS | PC2 (+) |
| WS_AREA | Upstream catchment area (watershed, log-transformed) | 30 – 43,496 | km<sup>2</sup> | --- | NHN | --- | PC3 (+) |

ARMOR = Global Observed Ocean Physics Reprocessing; AVISO+ = Archiving, Validation and Interpretation of Satellite Oceanographic data; GHCN = Global Historic Climate Network; MODIS = Moderate Resolution Imaging Spectroradiometer; NHN = National Hydrographic Network; ORAP = Global Ocean Physics Reanalysis ECMWF; PISCES = Global Ocean Biogeochemistry Non-assimilative Hindcast; WOA09 = World Oceanographic Atlas 2009

Candidate SNPs associated with environmental variables were identified using the complete genomic dataset with a combination of three methods: RDA, LFMM, and Baypass. Genes within 10,000 base pairs were recorded using bedtools (Quinlan, 2014) for candidate SNPs identified by at least 2 methods. We consulted the UniProt database (Uniprot Consortium, 2019) for information on biological gene functions in Atlantic Salmon, Zebrafish (*Danio rerio*), or other organisms.

#### 2.7.1 RDA

We used a redundancy analysis (RDA, Legendre & Gallagher, 2001) implemented in the R package *vegan* (Oksanen et al., 2012) to investigate multivariate correlations of genotypes (in the form of allele frequencies by population) with environmental variables. We took into account the structure in the data which derives from spatial patterns with a distance-based Moran’s eigenvector map (dbMEM; Borcard & Legendre, 2002; Peres-Neot & Legendre, 2010). To build this map, we transformed pairwise marine distances measured in the section above in Euclidian distances by creating a Delaunay graph with the function chooseCN in the R package *adegenet* (Jombart, 2008). We then used the R package *adespatial* (Dray et al., 2016) to compute the dbMEMs from the Euclidian distances. This approach was chosen over mere longitude and latitude information to avoid the underestimation of marine migration distances (e.g., by preventing considering impossible movements through landmass). Eigenvectors reflecting positive spatial autocorrelation were used as covariables in a partial RDA. As spatial structure could potentially hide evolutionary significant environmental gradients, we performed a second RDA excluding spatial eigenvectors correlated to environmental factors, as suggested by Forester, Lasky, Wagner and Urban (2018).

Spatial factors were first submitted to a backward model selection procedure with the function ordistep in *vegan*, and only significant (p < 0.05) covariables were kept. We then repeated this model selection method for environmental factors for both RDAs. To identify SNPs associated with environmental components, Z-scores were obtained for the distribution of individual SNP loadings on RDA axes explaining a significant portion of genetic variation (p < 0.05), and SNPs were defined as outliers if their maximal absolute Z-score was over 3.5 (p < 0.0002).

#### 2.7.2 LFMM

Latent factor mixed models (LFMM; Frichot, Schoville, Bouchard & François, 2013) determine loci-environment associations using a Bayesian mixed-model with environmental variables included as fixed effects. Latent factors are derived from a PCA and used as random effects to control for population structure. Missing data in the genetic dataset were imputed based on the most frequent genotype in the sampling site and we build the model using the lfmm_ridge function of the R package *lfmm* (Caye, Jumentier, Lepeule & François, 2019). The number of latent factors (K-value) and regularization parameters (lambda) were respectively set to 9 and 1e-5, to minimize predictor error estimated by a cross-validation method as advised in the *lfmm* manual. P-values were calibrated using the genomic control method and false discovery rate (q-value) was calculated following the Benjamini-Hochberg procedure in the *qvalue* R package. Associations between a SNP and environmental factor with q-value < 0.01 were considered significant.

### 2.7.3 Baypass

The standard covariate (STD) model implemented in Baypass assesses associations between allele counts and population-specific covariates using Bayesian Factors (BFs; Gautier, 2015). We estimated BFs by calculating the median of 5 runs of the STD model with the importance sampling (IS) algorithm (Coop, Witonsky, Di Rienzo & Pritchard, 2010). We controlled for the neutral population structure by providing the allele frequency covariance matrix (i.e., omega) that we computed using the core model implemented in Baypass using the same dataset. Default parameters were used for both the STD and core models. Pairs of markers and covariates with a median BF > 10 decibans (dB) were considered to have strong evidence for their association (Jeffreys, 1961, revised by Lee & Wagenmakers, 2013).

## 3. Results

Based on mtDNA, we only detected second contact between glacial lineages in Labrador: Samples from ANA and KAM mapped on previously published haplotypes from either the Atlantic (n = 23; ATL1, ATL4) or Arctic (n = 11; ARC19, ARC22) lineage. Mitochondrial haplotypes from Nunavik and Baffin Island samples all (n = 114) corresponded haplotypes from the Arctic lineage, primarily ARC19 (n = 85), but also ARC22, ARC25 and ARC26 (Table S1).

Using GBS, we sequenced a total of 745 samples, of which 95 did not pass our filtering criteria. Retained individuals (n = 650) had 2.217 million reads on average. A total of 30,753 SNPs were called and passed basic filtering in STACK2. Among these, 7,244 SNPs were categorized as duplicates and removed (Figure S1) and 5,009 SNPs were pruned during linkage disequilibrium assessment. We then identified and removed 408 SNPs linked to sex, 139 of which mapped on the sex chromosome (representing 15.6% of those SNPs) and 269 were elsewhere in the genome (1.5% of those SNPs). The final dataset used in subsequent analyses comprised 18,112 SNPs (see Table S2 for exact criteria and number of filtered SNPs), with a global 6.87% missing genotypes. Bayescan identified 273 SNPs as outliers (q-value < 0.05), 170 of those had higher F_ST_ values than expected under neutrality, suggesting divergent selection, and 103 had lower Fst values than expected, suggesting balancing selection. Pcadapt identified 186 outliers (q-value < 0.05), including 23 SNPs commonly identified by both genome scan methods, for a total of 436 markers putatively under selection. The remaining 17,676 SNPs were considered neutral for subsequent analyses.

### 3.1 Population structure

Population statistics are presented in Table 2. N_e_ ranged from 62 to 810 and was correlated with catchment area (30 – 43,496 km^2^, log-transformed, r = 0.46, p = 0.035), but the Payne River (PAY) had among the lowest N_e_ values (96) despite having the largest catchment area (when excluding PAY: r = 0.62, p = 0.004). The first axis of the PCoA reflected the longitude of the sampling sites, while the second axis differentiated populations in Western Ungava Bay from all other sampling sites (Figure 2).

**Table 2.** Summary of sampling and basic statistics. Samples were collected on adults in rivers and lakes, with exceptions († in estuaries, ‡ on juveniles).

| *Region* | *Code* | *Name* | *LON* | *LAT* | *n* | *H<sub>o</sub>* | *H<sub>e</sub>* | *Polymorphic*<br>*SNPs* | *N<sub>e</sub>*<br>*(IC 95%)* |
| --- | --- | --- | --- | --- | --- | --- | --- | --- | --- |
| *Nunavik* |  |  |  |  |  |  |  |  |  |
| *Hudson Bay* |  |  |  |  |  |  |  |  |  |
| *Southern* |  |  |  |  |  |  |  |  |  |
|  | IPI | Ipikutuk River | -78.38 | 58.73 | 24 | 0.272 | 0.254 | 9,748 | 195 (4.8) |
|  | SAP | Saputaliuk River | -78.36 | 58.7 | 17 | 0.292 | 0.260 | 9,519 | 62 (0.6) |
|  | FMI | Five Mile Inlet | -78.21 | 58.56 | 24 | 0.250 | 0.258 | 10,060 | 115 (1.3) |
| *Northern* |  |  |  |  |  |  |  |  |  |
|  | KOA | Korak River | -77.63 | 60.75 | 28 | 0.278 | 0.282 | 11,219 | 330 (8.1) |
|  | CHU | Chukotat River | -78.02 | 60.79 | 15 | 0.283 | 0.289 | 10,554 | 504 (34.9) |
|  | KOV | Kovik River | -77.7 | 61.36 | 21 | 0.267 | 0.282 | 10,704 | 78 (0.7) |
| *Hudson Strait* |  |  |  |  |  |  |  |  |  |
|  | BDE† | Déception Bay | -74.62 | 62.13 | 30 | 0.317 | 0.320 | 12,403 | 321 (6.1) |
|  | DUQ | Duquet Lake | -74.53 | 62.06 | 44 | 0.313 | 0.318 | 12,759 | 491 (8.8) |
|  | FRM | François-Malherbe Lake | -74.25 | 62.04 | 38 | 0.317 | 0.320 | 12,691 | 610 (15.9) |
|  | DOU | Douglas Harbour | -72.65 | 62.06 | 29 | 0.308 | 0.316 | 11,912 | 161 (1.7) |
| *Ungava Bay* |  |  |  |  |  |  |  |  |  |
| *Western* |  |  |  |  |  |  |  |  |  |
|  | PAY | Payne River | -70.7 | 60.01 | 30 | 0.300 | 0.311 | 11,791 | 99 (0.6) |
|  | CHR | Red Dog River | -69.79 | 59.3 | 39 | 0.291 | 0.294 | 11,500 | 172 (1.5) |
|  | VOL | Voltz River | -69.66 | 59.25 | 25 | 0.302 | 0.293 | 10,698 | 236 (4.6) |
|  | HAB† | Hopes Advance Bay | -69.63 | 59.32 | 24 | 0.296 | 0.298 | 10,864 | 215 (4.1) |
|  | FEU | Leaf River | -70.11 | 58.77 | 40 | 0.268 | 0.264 | 9,681 | 810 (31.7) |
|  | BER | Bérard River | -69.97 | 58.65 | 36 | 0.311 | 0.320 | 12,406 | 94 (0.4) |
| *Eastern* |  |  |  |  |  |  |  |  |  |
|  | GEO | George River | -65.95 | 58.69 | 37 | 0.313 | 0.314 | 12,036 | 787 (28.1) |
|  | AKI | Akilasaaluk River | -65.4 | 59.06 | 17 | 0.312 | 0.320 | 11,521 | 170 (3.5) |
| *Baffin Island* |  |  |  |  |  |  |  |  |  |
|  | AVA | Ava's Inlet | -72.64 | 64.01 | 23 | 0.285 | 0.289 | 11,148 | 254 (6.3) |
|  | LHO | Lake Harbour | -69.82 | 62.82 | 38 | 0.313 | 0.291 | 11,330 | 295 (4.4) |
|  | PRZ | Pritzler Harbour | -67.32 | 62.12 | 27 | 0.299 | 0.303 | 11,821 | 125 (1.2) |
| *Labrador* |  |  |  |  |  |  |  |  |  |
|  | KAM‡ | Kamanatsuk River | -62.54 | 56.74 | 18 | 0.310 | 0.307 | 11,656 | 303 (10.1) |
|  | ANA‡ | Anaktalik River | -62.15 | 56.49 | 26 | 0.302 | 0.300 | 12,011 | 81 (0.6) |

**Figure 2:**
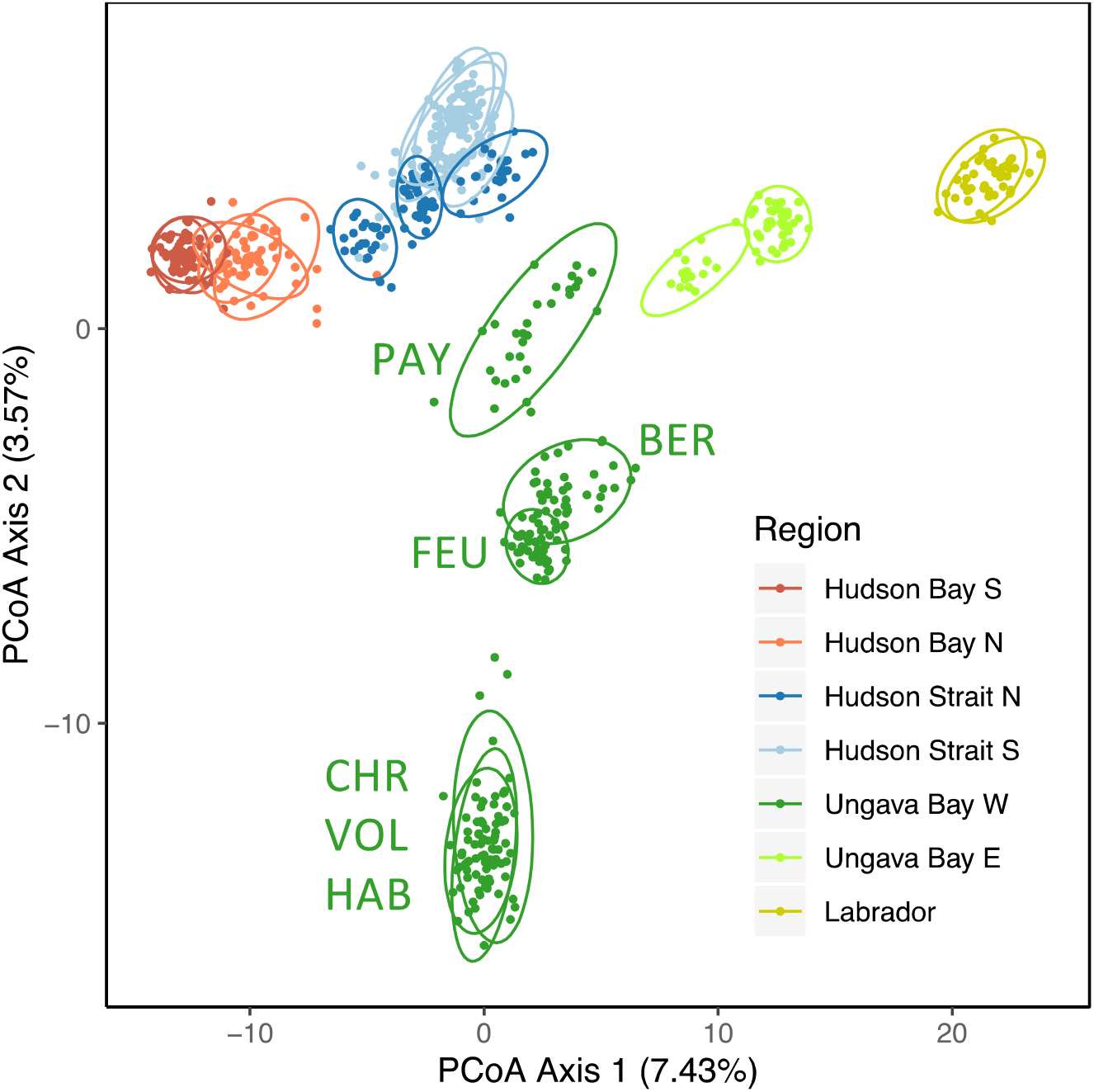
(A) Population structure assessed by a principal coordinate analysis (PCoA). Individual scores on PCoA axes 1 and 2 are presented as points and colored by a priori geographical region. An ellipse representing a 95% confidence interval was drawn around each sampling site. The percentage of genetic variance explained by each axis is in parentheses.

The number of genetic clusters best supported by an ADMIXTURE analysis with all non-estuarine sampling sites was 13 (Figure S2 for comparison of cross-validation errors). At this level, individuals within a sampling site were generally homogenous in their membership to clusters (Figure 3, see Figure S3 for K = 2 to 17). Sampling sites sharing a similar membership to clusters included sites in Labrador, two groups of three neighboring sampling sites in Northern and Southern Hudson Bay, and all pairs of rivers with a common estuary except the Leaf (FEU) and Bérard (BER) rivers. ADMIXTURE best supported K = 1 within those groups, but cross-validation errors for K = 2 were in some cases only slightly higher (Table S3). Individual cluster membership for K = 2 and K = 3 was then only loosely linked to the sampling site, except in Labrador sites, where samples were collected as juveniles.

**Figure 3:**
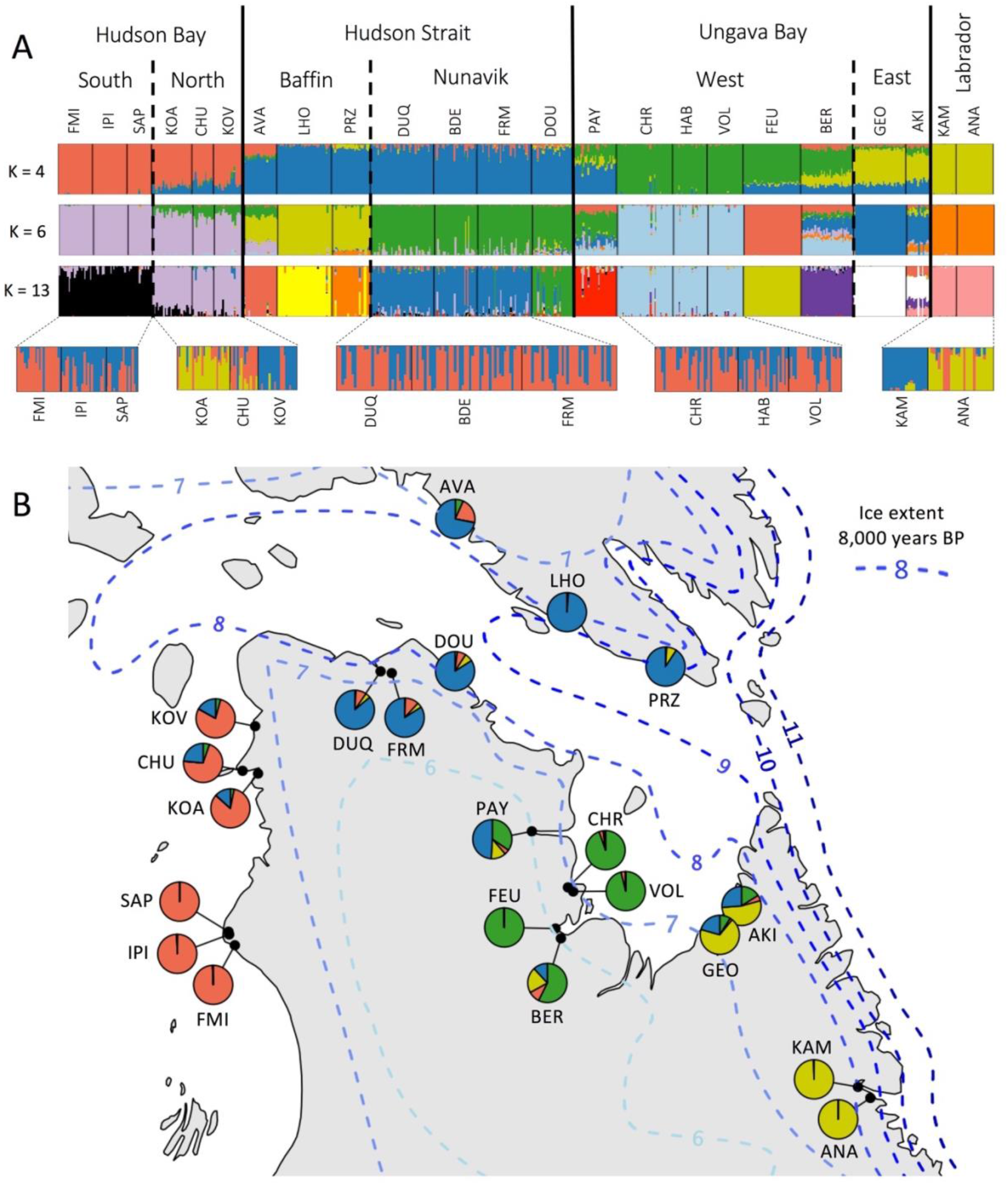
(A) Results of the hierarchical Bayesian clustering analysis implemented in ADMIXTURE for a number of genetic clusters (K) of 4, 6 and 13. Lower rows display the results for separate analyses on sampling sites sharing similar membership to clusters at K = 13, which yielded the lowest cross-validation error (CV). (B) Results of ADMIXTURE for K = 4 clusters, with individual ancestry averaged by sampling site and represented by pie charts. Approximative extent of glaciers, adapted from Dyke (2004), are represented by blue dashed lines for 6,000 - 11,000 years before present (BP).

Individual mean proportion of heterozygous markers varied regionally (Figure 4). Notably, southern Hudson Bay displayed the lowest observed proportion of heterozygous markers, and both southern and northern Hudson Bay had lower values than Hudson Strait (p < 0.05). In Ungava Bay, eastern sampling sites had a diversity similar to Labrador, and higher diversity than western sampling sites.

**Figure 4:**
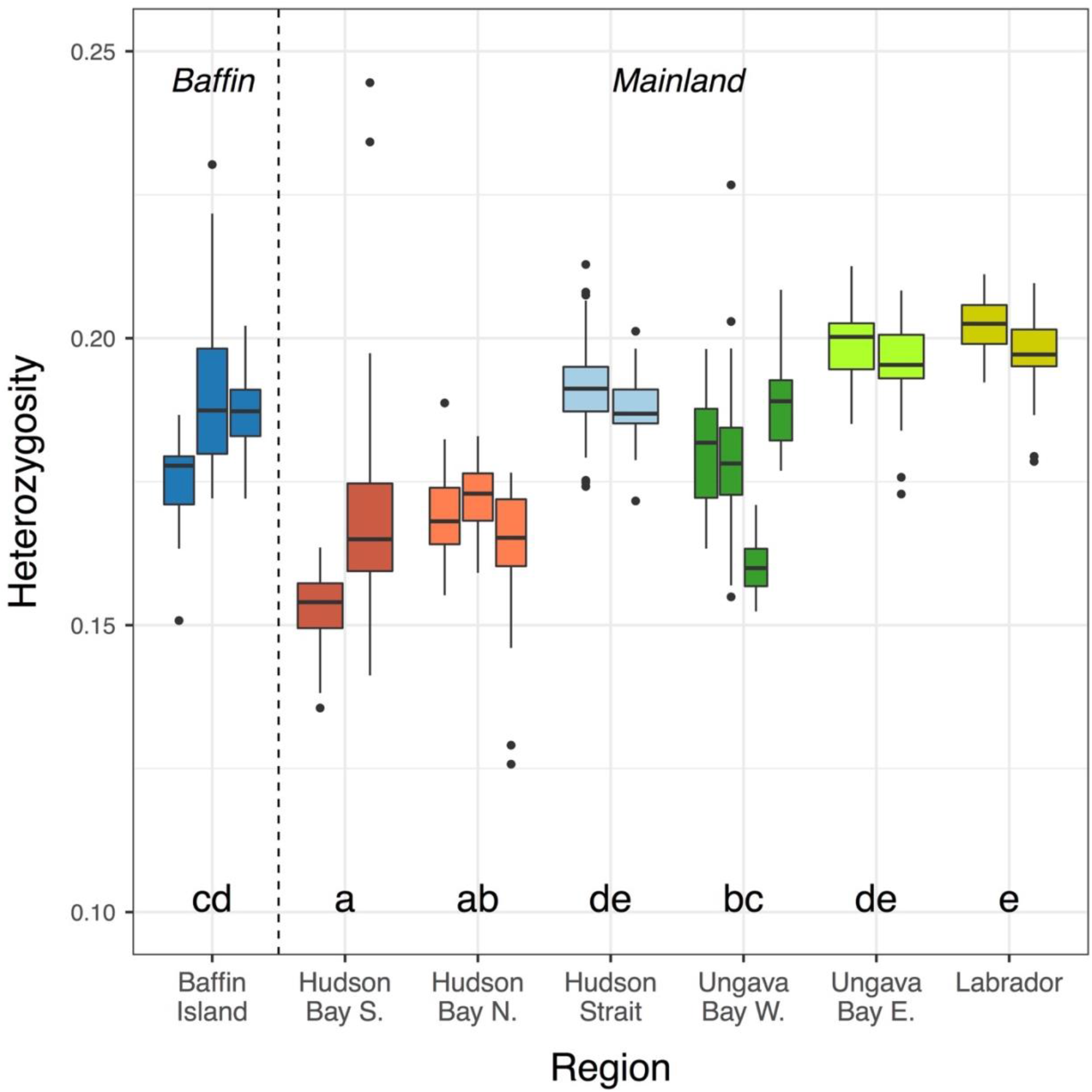
Individual proportion of heterozygous SNP markers in sampling sites, ordered following the coast from west to east, and colored by region. Sampling sites on connecting rivers and estuaries were grouped, except for FEU and BER. For each boxplot, bold line indicates mean, the box limits 25^th^ and 75^th^ percentile, and whiskers represent 10^th^ and 90^th^ percentile. Letters indicate group membership based on a comparison of least square means in a mixed-effect model (p < 0.05).

### 3.2 Landscape genomics

F_ST_ calculated by pairs of sampling sites ranged between 0.000 and 0.319 across neutral loci and between −0.003 and 0.544 across putatively adaptative loci (Figure 5A, Table S4). F_ST_ for adaptive loci were roughly twice the value for neutral loci and both measures were highly correlated (slope = 1.78, r = 0.94). All p-values for comparisons using neutral loci were under 0.01, except for a pair of sampling sites on rivers sharing an estuary (p_CHR-VOL_ = 0.568) and a pair of sampling sites on a river and its estuary (p_BDE-FRM_ = 0.015). These results suggest that all unconnected systems likely harbour genetically distinct populations, although their extent of differentiation is highly variable.

**Figure 5:**
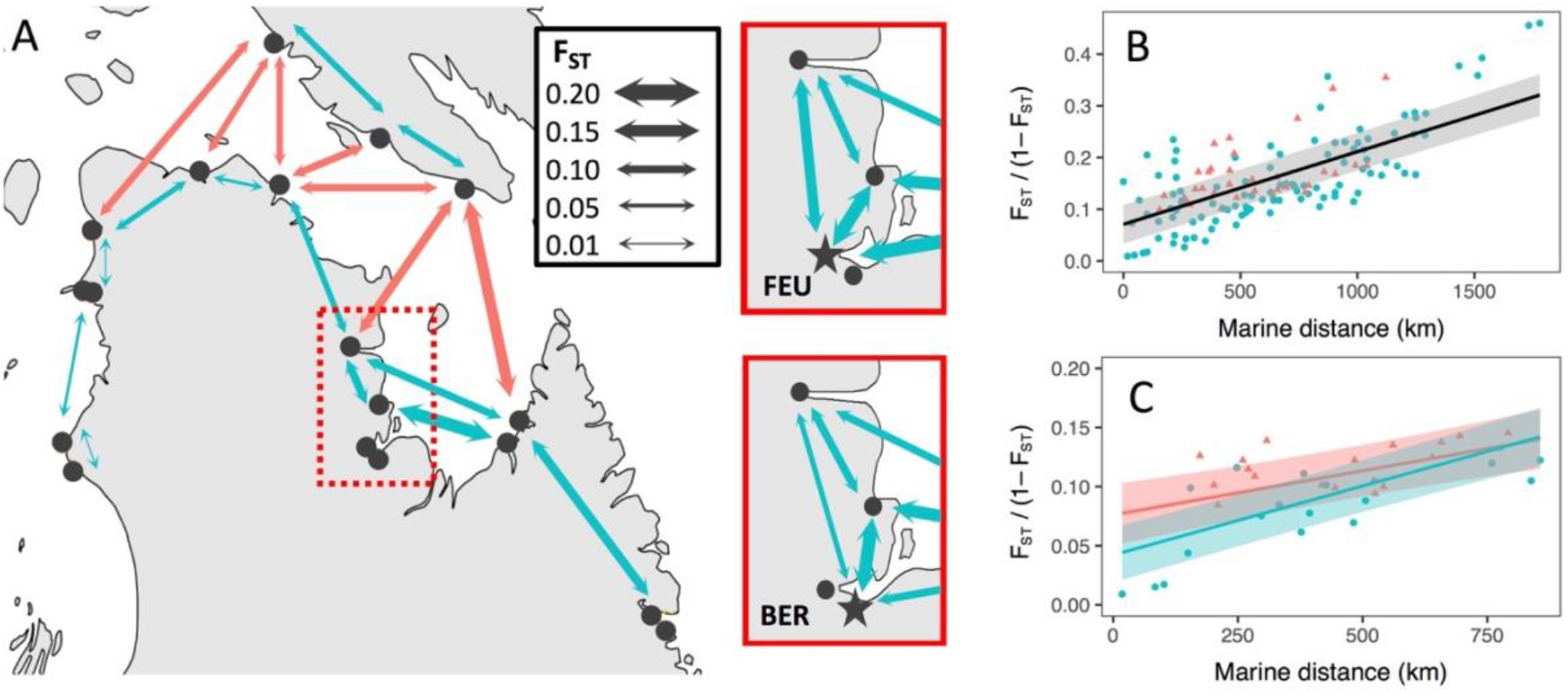
(A) Pairwise F_ST_ between neighboring sampling sites are shown, with line thickness proportional to values. All pairwise F_ST_ values are presented in Table S3. Red extent is magnified to contrast F_ST_ between star-shaped Leaf (FEU) and Bérard (BER) rivers (pairwise F_ST_ = 0.135). Other pairs of adjacent sampling sites had their F_ST_ values averaged for better visualization. (B-C) Isolation-by-distance represented by relation between marine distances and linearized pairwise F_ST_, estimated between pairs of sampling sites separated by Hudson Strait (red triangles) and on the same coast (blue dots). Regression lines and 95% confidence intervals are plotted for mixed-effect models with fixed effects including marine distance, crossing of Hudson Strait and the interaction of those factors. Models were fitted using (B) all sampling sites and (B) only sampling sites within 250 km of Hudson Strait.

When examining IBD over all sampling sites, a positive correlation was found between linearized F_ST_ and marine distances (Figure 5B), but the inclusion of the effect of crossing the Hudson Bay (CROSSHS) did not improve the model, based on marginal R^2^ (Table 3). However, similar analyses focusing on sampling sites within 250 km of the Hudson Strait showed higher linearized F_ST_ in pairs of populations on either side of the Hudson Strait than in pairs on the same coast (Figure 5C).

**Table 3.** Parameters of isolation-by-distance mixed effect models, displaying degrees of freedom (df), conditional Akaike information criteria (cAIC) and marginal R-Squared (R^2^m) compared to the null model.

|  | All sites |  |  | Sites within 250 km of Hudson Strait |  |  |
| --- | --- | --- | --- | --- | --- | --- |
| | df | cAIC | $R^2m$ | df | cAIC | $R^2m$ |
| FST ~ 0 | 17.18 | -362.18 | -- | 8.06 | -144.37 | -- |
| FST ~ DIST | 18.95 | -553.38 | 0.655 | 10.62 | -199.86 | 0.535 |
| FST ~ DIST + CROSSHS | 19.80 | -553.02 | 0.651 | 11.65 | -211.28 | 0.626 |
| FST ~ DIST * CROSSHS | 20.80 | -552.02 | 0.649 | 12.64 | -214.37 | 0.642 |

### 3.3 Gene-Environment Association

Environmental factors were each summarized with 3 PC axes (Table 1). The first axis (PC1; 38.0% of variation) was associated to primary productivity, dissolved oxygen, precipitation and winter air temperature. The second axis (PC2; 26.5% of variation) was associated to summer temperature, both in the marine and freshwater habitats, as well as to turbidity. Finally, the third axis (PC3; 17.2% of variation) related mostly to tidal amplitude and partly to size of the catchment area (Figure S4, S5).

The distance-based Moran’s eigenvector map produced 12 eigenvectors reflecting negative autocorrelation and 4 eigenvectors for positive autocorrelation (MEM1 – 4, see Figure S6). Model selection for the RDAs excluded MEM3 (p = 0.13), MEM4 (p = 0.21). MEM1 was highly correlated with PC3 (r = 0.71) and MEM2 was linked to PC1 (r = 0.74). Thus, no spatial covariable was included in the RDA excluding spatial factors correlated to the environment, which we further simply refer to “RDA”, compared to the partial RDA (thereafter “pRDA” and including MEM1 and MEM2 as covariables). Spatial covariables explained 34.8% of the genetic variance, 29.0% of which was shared with environmental factors, resulting in the RDA and pRDA having respective adjusted R^2^ of 0.438 and 0.148.

In both the pRDA and the RDA, the three first axes explained significant portions of the variance (p < 0.05) and were used for outlier detection. The pRDA identified a total of 295 outlier SNP markers distributed across 36 Arctic Char linkage groups (Figure S7). Respectively 98, 113, and 84 outliers were found on each of the three axes (Figure 6). Those outliers were generally most correlated to PC1 (n = 185) or PC2 (n = 90). The RDA identified only 14 outliers on the first axis, and respectively 112 and 109 on the second and third axes (Figure S8), for a total of 234 markers on 38 linkage groups (Figure S9). The environmental components PC1, PC2, and PC3 had respectively 62, 83, and 90 of those outliers most correlated to them. A total of 55 markers were identified as outliers by both the pRDA and the RDA, most of which were most correlated to PC2 (n = 35).

**Figure 6:**
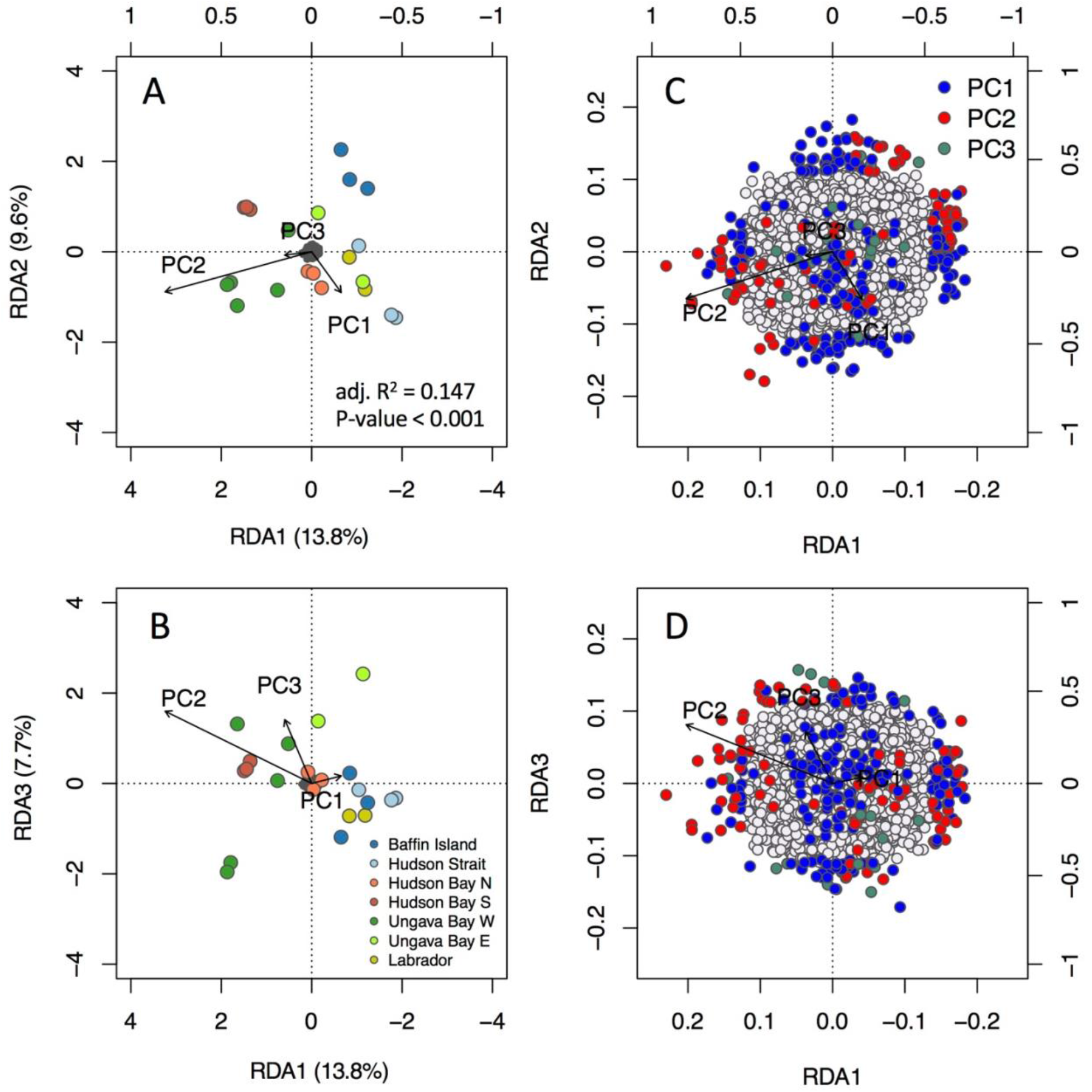
Triplots for (A) axes 1 and 2, and (B) axes 1 and 3 in a pRDA including spatial components. The dark grey cloud of points at the center of each plot represents the SNPs, and coloured points represent sampling sites with color coding by region. Triplots are magnified to highlight SNP loadings on (C) RDA axes 1 and 2, and (D) axes 1 and 3. Candidate SNPs are shown as colored points with coding by most correlated environmental predictor (see text for description of predictors). Vectors represent environmental predictors, according to the scales on top and right axes.

LFMM identified a total of 173 outlier SNPs (q < 0.01, Figure S10), 13 of which being associated PC1, 102 associated to PC2, and 58 to PC3. A total of six SNPs were significantly associated with more than one environmental component, five of which were associated to both PC2 and PC3. The IS model under Baypass identified 55 SNPs (BF > 10 dB, Figure S11). Of these, 22 were associated to PC1, 13 to PC2, and 20 to PC3.

Of the 613 unique outliers, 75 (12.2%) were identified as associated with an environmental factor by at least two GEA methods (Figure 7, see Figure S12 for distribution of allele frequencies). Among those, 21 were considered associated to different factors depending on the method. Each of those mismatches included the RDA/pRDA method, for which the candidate SNPs were most correlated to a certain factor, but also had an absolute correlation coefficient over 0.45 for the factor identified by the other method (LFMM or BAYPASS). As such, we reported only the environmental factors detected by LFMM or BAYPASS in such cases. Thus, six candidate SNPs were associated to PC1 (precipitation, winter air temperature and productivity), 53 were associated to PC2 (air and sea-surface summer temperature, salinity, and turbidity) and 16 were associated to PC3 (tides). We identified a total of 67 named genes within 10,000 base pairs of those candidate SNPs, covering a wide range of biological functions, including muscle contraction, development, and circadian rhythm (Table S7).

**Figure 7:**
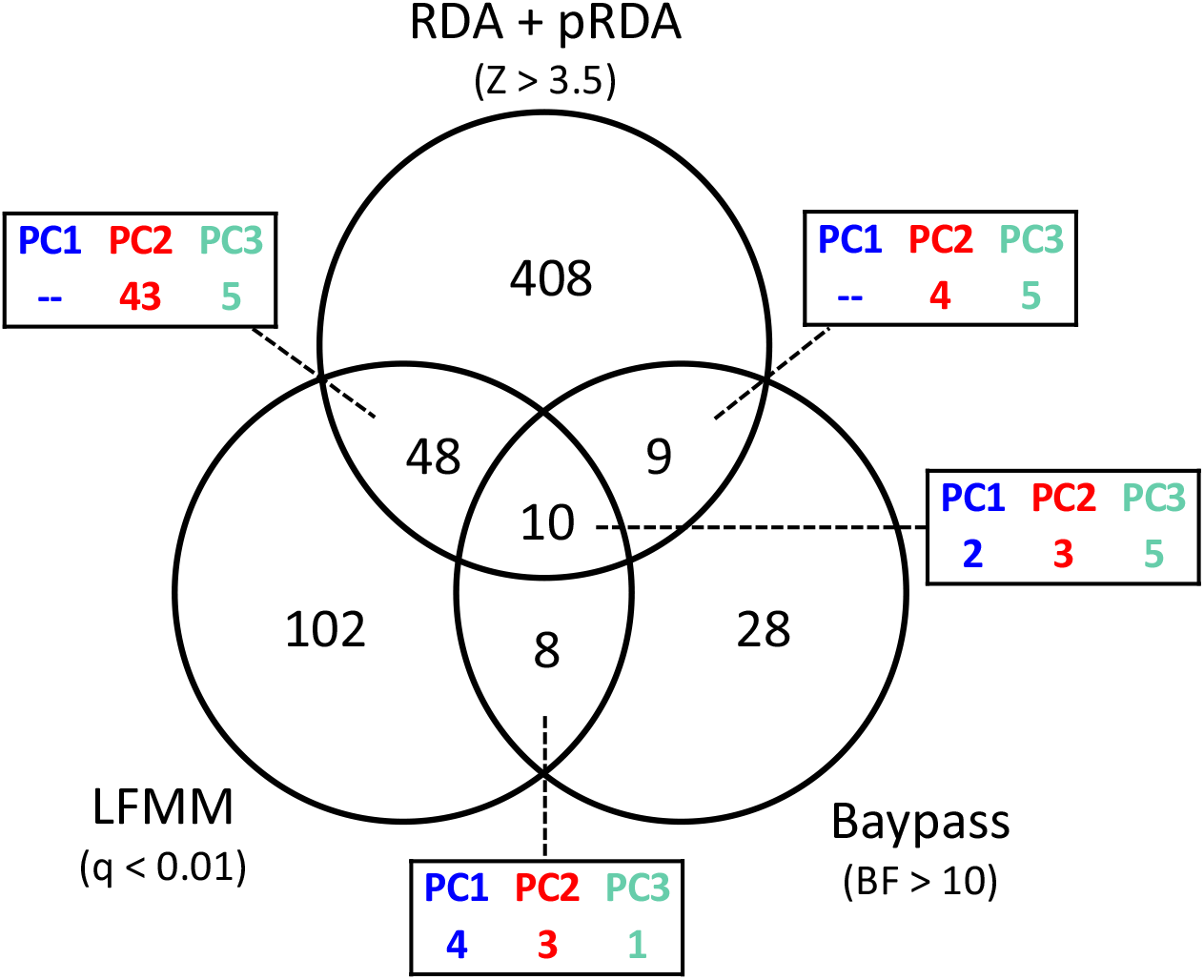
Intersection of candidate SNPs detected by three GEA methods. Candidates from the redundancy analyses with (pRDA) and without (RDA) spatial correction were combined. For each intersection, the box indicates the distribution of environmental components associated to candidate SNPs.

## 4. Discussion

Anadromous salmonids are considered prime models for understanding local adaptation (Fraser et al, 2011) and the many landscape genomics studies which they have been the subject of have provided key information for the conservation and management of economically important species (Waples, Naish & Primmer, 2020). Here, we documented for the first time patterns of neutral and adaptive genetic variation in anadromous Arctic Char populations in the Nunavik region using genomic data. By combining fine- and broad-scale sampling, our results revealed a hierarchical genetic structure. Neighboring hydrographic systems generally harbored distinct populations that could be regrouped within oceanographic basins, while differentiation was also influenced by isolation-by-distance. To assess the potential for local adaptation to the contrasted environments in our study area, we combined three GEA methods. By doing so, we found genomic signatures that were consistent with local adaptation to either freshwater or marine habitats.

### 4.1 Neutral structure in a post-glacial context

By 8,000 YBP, most of Hudson Strait and the coast of Labrador was free of ice while most of the Hudson Bay and the Ungava Bay remained glaciated. By 6,500 – 7,200 YBP, the coasts of Hudson Bay and Ungava Bay deglaciated, meaning those regions would be the last to have been colonized by Arctic Char. The decreasing genetic diversity we observed along the coast from Hudson Strait to southern Hudson Bay, as well as low F_ST_ values between populations within the Hudson Bay region, are therefore consistent with expectations of a recent range expansion (Eckert, Samis & Lougheed, 2008; Goodsman, Cooke, Coltman & Lewis, 2014). Similar patterns were found in European lamprey species (*Lampetra* spp.), with diversity and differentiation decreasing in populations far from the Iberic glacial refugia (Mateus et al., 2018), and in Scottish populations of Atlantic Salmon, where genetic diversity was lower in more recently deglaciated regions (Cauwelier et al., 2018).

In Northern Labrador, Salisbury et al. (2019) found extensive admixture of the Arctic mitochondrial lineage, which crossed the Canadian Arctic Archipelago from a glacial refugium in the western Arctic (Brunner et al., 2001; Moore et al., 2015), and the Atlantic lineage, which we expect to have crossed the Atlantic Ocean from the Palearctic during the last deglaciation (Brunner et al., 2001; Wilson, Hebert, Reist & Dempson, 1996). Our results were consistent with these previous studies as we did not observe any Atlantic lineage haplotypes in populations outside Labrador. However, similarly to Moore et al. (2015), we found signs of introgression from the Atlantic lineage in our nuclear data from southern Baffin Island, and even stronger evidence in Eastern Ungava Bay, where our ADMIXTURE results showed shared ancestry with Labrador samples. Such mito-nuclear discordance is not uncommon in cases of sex-biased dispersion and adaptive introgression (Toews & Brelsford, 2012), which we did not investigate here. Nevertheless, admixture in the eastern part of our study area could explain the higher genetic diversity, as expected during secondary contact of marine fishes (Bay & Caley, 2011; Grant & Bowen, 1998).

### 4.2 Contemporary gene flow

Our results revealed a hierarchical genetic structure with most geographic regions containing distinct populations of Arctic Char in every sampled river not sharing an estuary. However, population structure in Hudson Bay appeared weaker, as there were signs of admixture between rivers within 100 km of each other. Other studies on Arctic Char have attributed low genetic differentiation over long distances to recent post-glaciation colonization events (Moore et al., 2013; O’Malley et al., 2019). However, lower salinity and a longer summer period in Hudson Bay could also lead to higher connectivity between estuaries.

As migration in Arctic Char has been found to be predominantly coastal (Moore, 1975; Moore et al., 2013, 2015; Spares et al., 2012), we expected the Hudson Strait, a 120-km-wide open water area, to restrict gene flow between populations on opposite shores. However, those pairs of populations did not deviate from IBD, which could suggest ongoing gene flow over long marine stretches. In contrast, we did find limited evidence for Hudson Strait acting as a barrier to gene flow at a finer spatial scale, i.e. by restricting analyses to sampling sites around the strait. Differentiation between populations separated by the Hudson Strait was lower than, for example, pairwise F_ST_ between populations in western and eastern Ungava Bay, despite those populations being connected by a near-shore migration route. We cannot exclude the possibility of other barriers to gene flow in our study area, as near-shore oceanographic features could also limit dispersal (e.g., Quéméré et al., 2015), but differentiation in Ungava Bay could also be driven by nuclear introgression from the Atlantic lineage in the eastern sampling sites. Additionally, populations on either side of Hudson Strait could share ancestral polymorphism rather than exhibit contemporary gene flow. As this system is unlikely to be at migration-drift equilibrium due to its young evolutionary age, these other processes (introgression and ancestral shared polymorphism) could hide the effect of the Hudson Strait as a potential barrier to gene flow. In this way, our results hint again at the relative importance of glacial lineages over contemporary restrictions to gene flow in this recently recolonized area.

Arctic Char is known for its higher straying rates than other salmonids, but many studies argue that this dispersal does not necessarily lead to gene flow, as individuals are more prone to straying when overwintering than when breeding (Moore et al., 2013; Moore et al. 2017; Sévigny, Harris, Normandeau, Cayuela, Gilbert & Moore, in prep). In this study, adults sampled in rivers sharing an estuary generally did not exhibit population structure, suggesting high levels of straying between nearby rivers. However, in a more surprising case, there was no signs of adult dispersers between the Leaf River (FEU) and the Bérard River (BER), which also share an estuary. Interestingly, those two rivers are on either side of the limit of two major geological provinces (Thériaut & Beauséjour, 2012). As salmonids rely partly on their olfaction to recognize their natal rivers (Keefer & Caudill, 2014), the distinct geologies of the Leaf and Bérard systems could perhaps improve their homing ability and thus limit straying in these systems.

### 4.3 Genomic evidence for local adaptation

We explored how genetic variation in Arctic Char was linked to a range of climatic and physical environmental predictors while considering both freshwater and marine habitats, something that has rarely been done in salmonids (but see Bekkevold et al., 2019). While different GEA methods identified candidate SNPs with every environmental component considered, we found that most of the best-supported candidates were associated with the component reflecting summer SST and air temperature, salinity, and turbidity (PC2).

In our study, we treated the environment experienced by anadromous Arctic Char over their life cycle as a complex set of collinear variables, hence leading to our inability to differentiate signatures of selection in the freshwater and marine habitats. We also created two sets of habitat-specific environmental components, but marine and freshwater components were strongly correlated and created several instances of candidate SNPs associated to both a marine and freshwater component by GEA methods (results not shown). Nevertheless, using those habitat-specific components, we did find through a constrained ordination that both habitats independently explained significant proportions of the genetic variation. For instance, measures of air temperature (as a surrogate for river temperatures) and sea temperatures were highly correlated, but likely caused selective pressures at different life stages. Several studies have found temperature to be a driver of genetic structure and local adaptation in salmonids, with most studies focusing on freshwater measures (Dionne et al., 2008; Bourret et al., 2013; Perrier et al. 2017; Silvester et al. 2018). In contrast, selective pressures in marine habitats have been argued to be weaker since observed mortality rates are lower at sea (Garcia de Leaniz et al., 2007; Quinn, 2005). However, SST near the mouth of spawning river was found to be correlated with timing of migration in species of Pacific salmon (Kovach, Ellison, Pyare & Tallmon, 2015) and to phenology-related genes in Arctic Char (Madsen et al., 2019). Sampling sites across our study area displayed considerably different SST conditions: while coastal waters surrounding sampling sites in Hudson Bay reached mean summer SSTs ranging from 5.5 – 7.5°C, Hudson Strait stayed closer to the freezing point (0.5 – 2°C). Such contrasts in surface temperature, coupled with a discrepancy in tidal regimes and salinity, likely produce widely different coastal habitats, which we argue could result in local adaptation of Arctic Char populations.

Recent studies of local adaptation in salmonids have focused on tributary-specific variation in freshwater conditions within a single catchment, (e.g., the Columbia River; Hand et al., 2016; Hecht et al., 2015, Micheletti et al., 2018). However, the freshwater environmental factors used in our study are catchment-based given our sampling strategy., which prevents us from knowing the precise spawning site or overwintering lake. In marine systems, genomic evidence for local adaptation and isolation-by-environment has been found both at local scales in heterogeneous habitats (e.g., Lenhert et al., 2019; Miller et al., 2019) and over large geographic distances (e.g., Clucas, Lou, Therkildsen & Kovach, 2019). Arctic Char is expected to use preferred habitats based on temperature, salinity, and prey availability (Harris et al., 2020; Spares et al., 2012; 2015). As we averaged near-shore marine conditions around river mouths, this study is limited to broad-scale environmental heterogeneity. This is in line with the suggestion by Fraser et al. (2011), that adaptation of anadromous salmonids to the marine environment should occur at a larger spatial scale than in fresh water.

Regardless of the geographic scale being studied, an ideal sampling design for detecting local adaptation should maximize the environmental variation while minimizing its collinearity with neutral genetic patterns. For example, Lotterhos and Whitlock (2015) suggested sampling pairs of populations with similar ancestry and contrasting environments. Such a design is suitable when studying variables that may change drastically over short distances, such as catchment area and upstream migration distances. However, climatic and physicochemical conditions experienced by geographically close populations of Arctic Char are more likely to be similar than in distant ones. Similar considerations were discussed in Nadeau, Meirmans, Aitken, Ritland and Isabel (2016), where patterns of isolation-by-distance, isolation-by-environment, and isolation-by-colonization were difficult to disentangle for two pine species in a recently recolonized range. In our study, some genotypes for GEA candidates (Figure S12) have varying allele frequencies that follow spatial patterns reminiscent of the neutral structure we described earlier, especially in PC1, where environmental variation followed a longitudinal gradient. In some of those cases, it is possible that a GEA was detected even though neutral processes could better explain the distribution of the observed allele frequencies. For example, if colonization of Hudson Bay did occur by rapid demographic expansion as discussed earlier, allele-surfing events, where a mutation can reach high frequency by chance alone (Edmonds, Lile & Cavalli, 2004), could explain differential allele frequencies (Rougemont et al., 2020). Alternatively, the introgression of the Atlantic glacial lineage in the eastern part of the study area could also have led to divergence due to drift during the LGM to be falsely identified as linked to environmental variation. On that note, the pRDA, LFMM, and Baypass methods all accounted in some way for neutral or spatial structure. However, it is noteworthy that even the RDA, while excluding spatial correction, only identified a few outliers whose variation was driven by the Labrador sampling sites, i.e. from the Atlantic glacial lineage. As we found a significant part of genetic variation to be explained jointly by environmental variation and spatial patterns, we advocate that including GEA methods that do not account for spatial structure offer a way to acknowledge that local adaptation can also contribute to genetic structure across contrasting environments, e.g., by selecting against maladapted migrants (Dionne et al., 2008; Wang & Barburd, 2014). In that sense, local adaptation in the system studied here could reinforce the neutral structure discussed earlier.

### 4.4 Implications for conservation and management

Genomic data has the potential to improve the definition of management and conservation units by accounting for neutral genetic structure as well as local adaptation (Bernatchez et al., 2017; Funk et al., 2012). Arctic Char populations in Nunavik support important small-scale subsistence fisheries, and stocks are managed on a river-by-river basis on the premise that each river contains a single and distinct population (Johnson, 1980). However, the very low genetic differentiation in rivers sharing an estuary and our inability to identify substructure at this level suggest high levels of straying in adults, in line with the evidence for the prevalence of mixed stocks of Arctic Char in adjacent rivers (Boguski, Gallagher, Howland & Harris, 2016; Moore, Lewis & Tallman, 2014; Moore et al., 2017). If genomic tools were to be used for stock assignment of adult fish (e.g., Meek et al., 2016; Moore et al. 2017), further local sampling including juveniles and/or reproducing adults would be useful to better understand fine-scale population structure. The evidence we found for local adaptation to environments is in concordance with the neutral genetic structure at a broader scale, as major oceanographic basins around Nunavik are contrasted both in their environments and ancestry of their Arctic Char populations. We therefore suggest that these regional differences between Hudson Bay, Hudson Strait, and Ungava Bay (with distinction of the western and eastern coast) should form the basis of management actions at a regional level.

Also, there is growing interest in Arctic Char hatchery projects in Nunavik, both for supplementation and reintroduction of Arctic Char in traditional fishing locations (George, 2007; Rogers, 2015). The genetic information gathered here could be of great use for those initiatives, and the adaptative variation explored in this study highlights the need for careful choices of source populations for broodstocks, as maladapted domesticated individuals can waste efforts and resources, in addition of likely being detrimental to wild populations (Fraser, Minto, Calvert, Eddington & Hutchings, 2010; Tymchuk, Biagi, Withler & Devlin, 2005).

As the Arctic warms at a greater pace than any other regions on earth (Cohen et al., 2014), there might be concern about the response of Arctic Char populations to their changing environment. Traits that are currently optimally adaptive in the present environment could eventually become maladaptive. Species will thus likely need to shift their distributions poleward and/or will need to adapt in order to persist in their current distribution if there is presence of appropriate genetic diversity/phenotypic plasticity. A temporal study recently showed that Arctic Char populations in Greenland have exhibited stable genetic structure over the last 60 years in face of rapid climate change, and argued that gene flow, although low, could allow for a modest level of evolutionary rescue in the short term (Christensen, Jacobsen, Nygaard & Hansen, 2018). Our study shows potential for local adaptation of Arctic Char populations to both their marine and freshwater habitats. As changes in climate might operate at a different pace, scale, and stability in marine and terrestrial ecosystems (Burrows et al., 2011), there is a need for continued research about the interaction of selective pressures over the lifespan of anadromous organisms.

## Data availability Statement

Raw sequences that support the findings of this study are openly available on NCBI SRA, accession number PRJNA655216 (https://www.ncbi.nlm.nih.gov/bioproject/PRJNA655216/).

