## Supplemental Figures for "Population structure and genomic evidence for local adaptation to freshwater and marine environments in anadromous Arctic Char (*Salvelinus alpinus*) throughout Nunavik, Québec, Canada": SAAL_Nunavik_EvoApp_SUPPfig_26nov.pdf

Figure S1: Relationship between the allelic ratio (MedRatio), the proportion of heterozygotes (PropHet), the proportion of rare homozygotes (PropHomRare) and FIS for 31,535 SNPs after basic filtering in Stacks 2. SNPs were categorized in duplicates (red), diverged duplicates (blue), high coverage (green), low confidence (purple), or insufficient number of samples with the minor allele (orange), based on criteria in Table S1

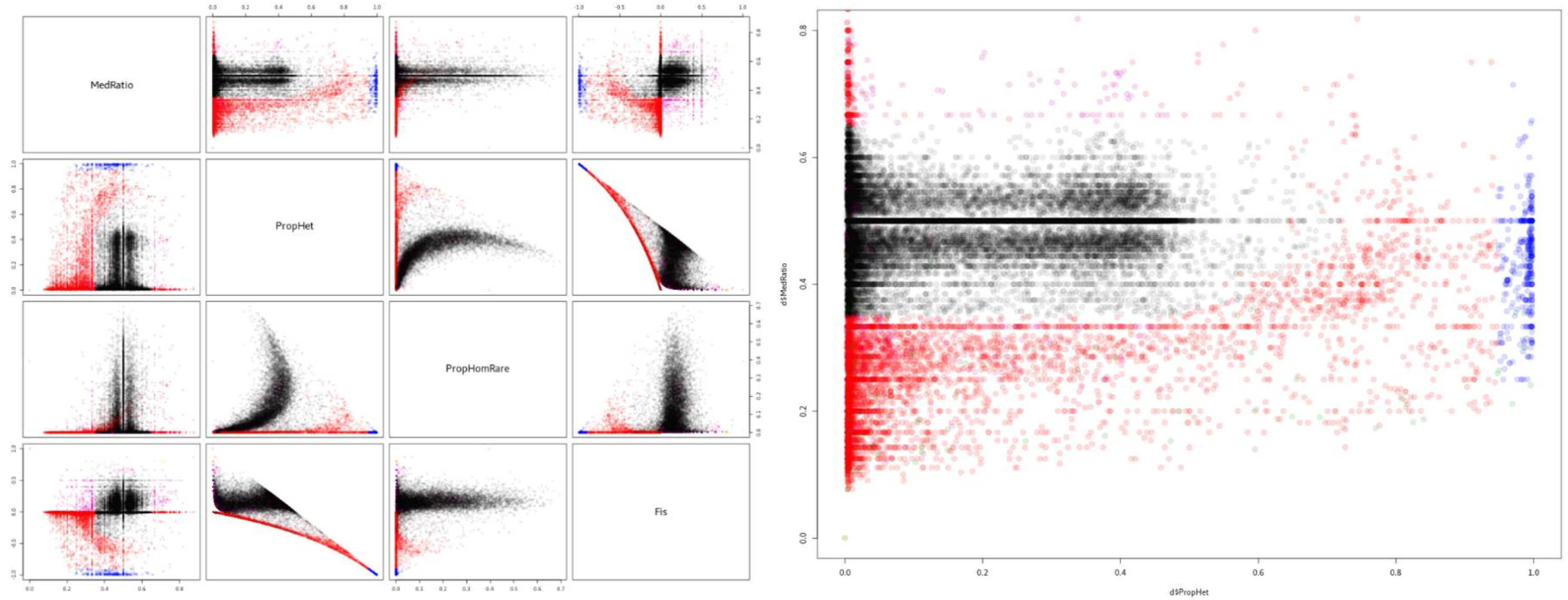

Figure S2: Cross-validation (CV) error for an ADMIXTURE clustering analysis according to the number of considered clusters (K), varying from K = 1 to 20.

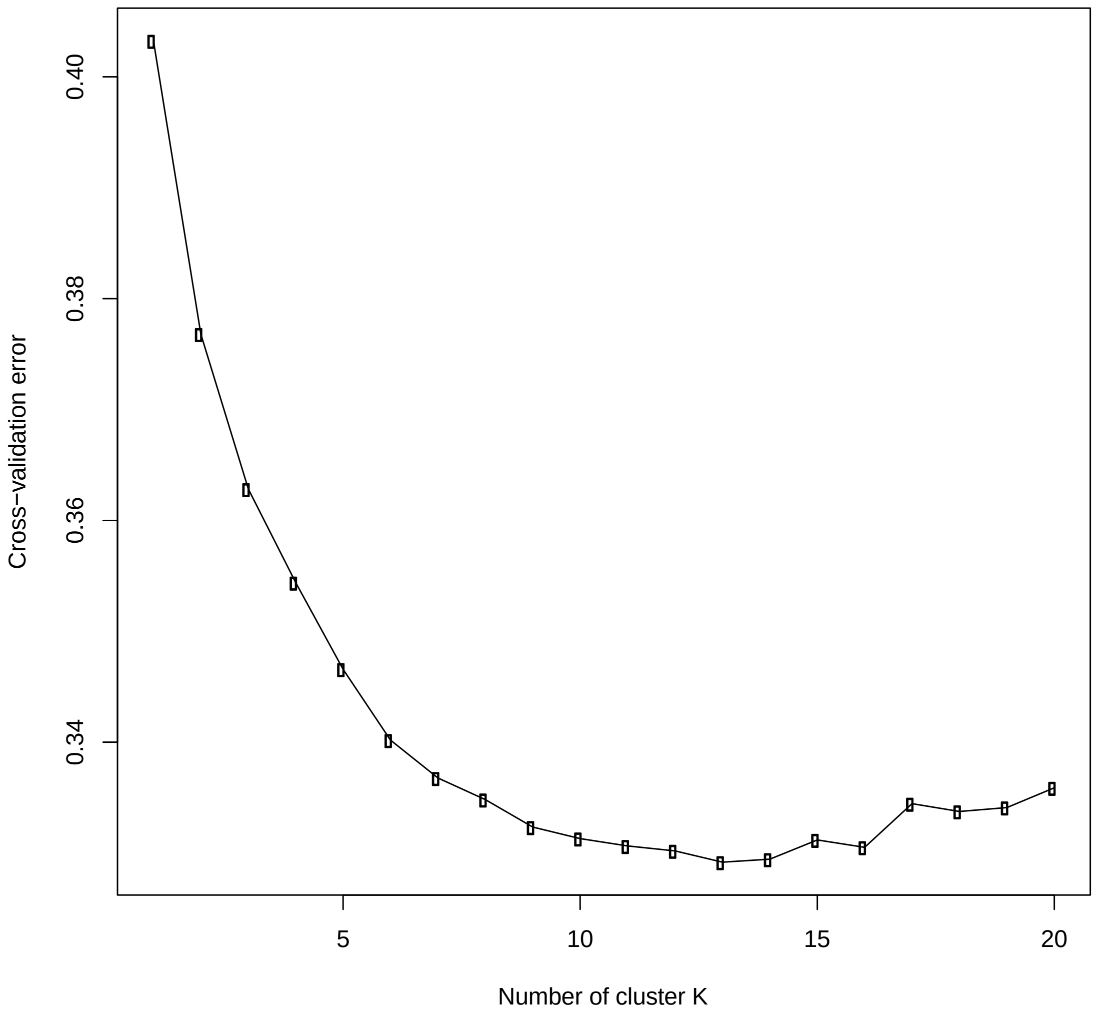

Figure S3: Results of the hierarchical Bayesian clustering analysis implemented in ADMIXTURE for a number of genetic clusters (K) ranging from 2 to 16.

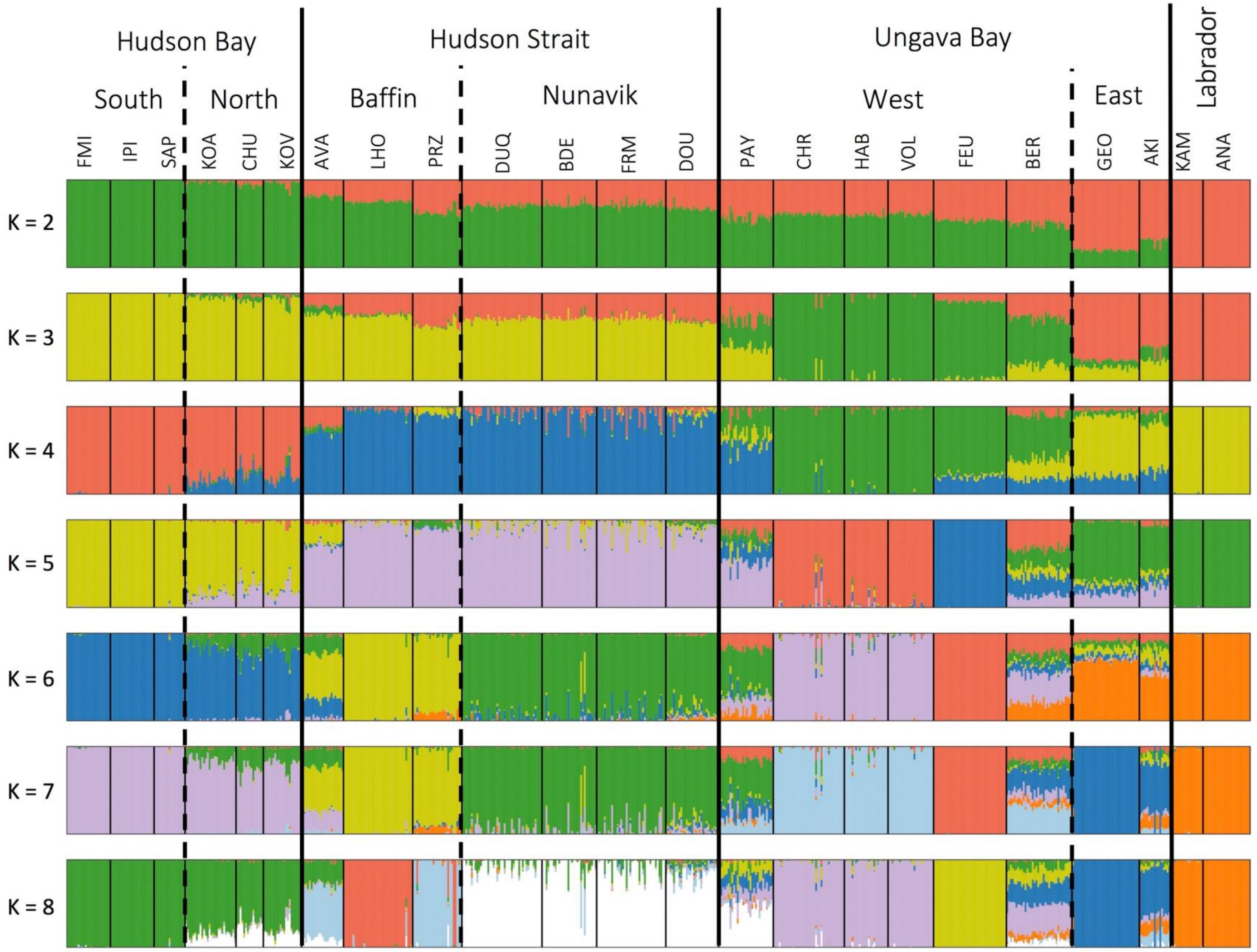

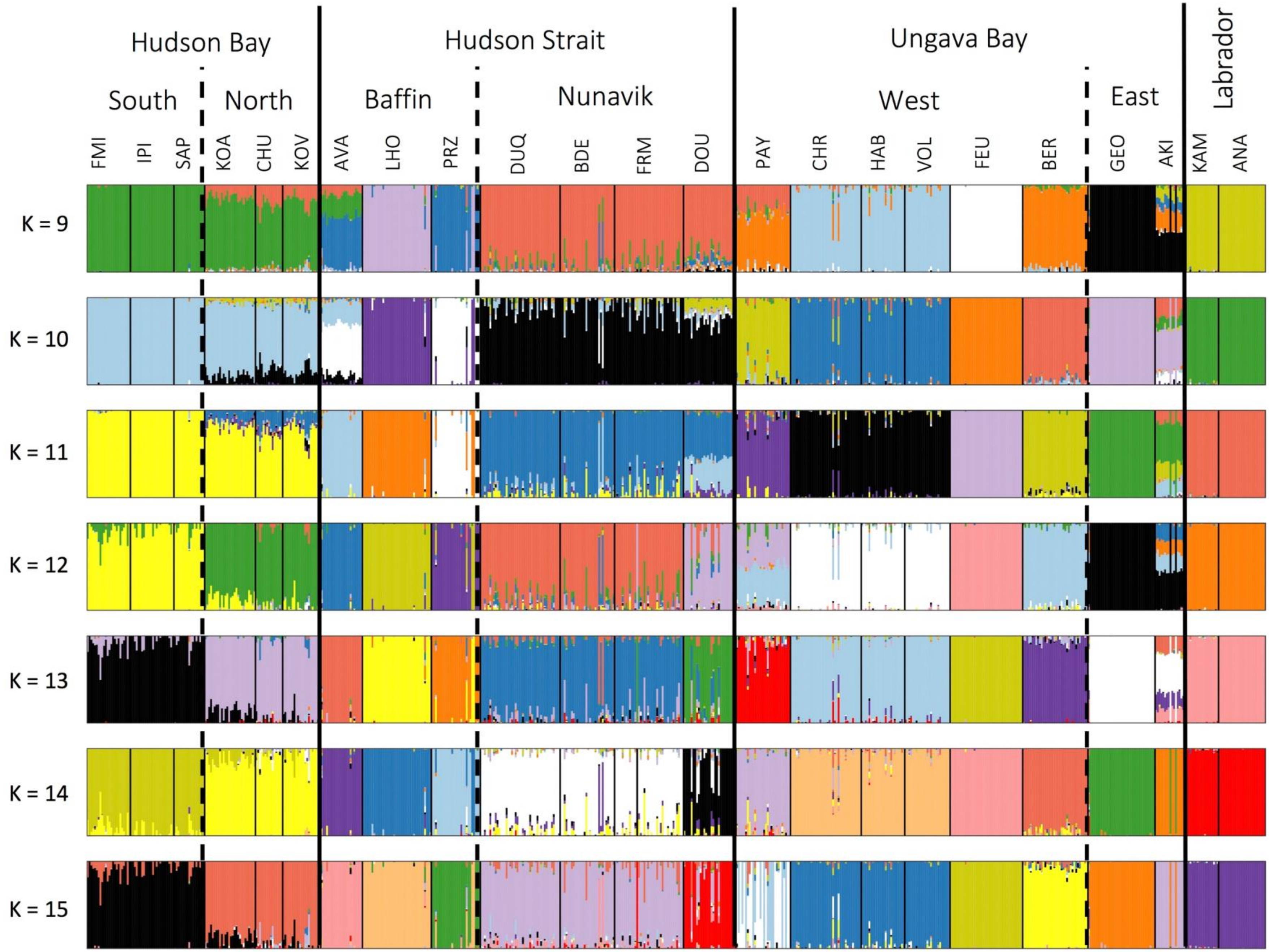

Hudson Bay
Hudson Strait
Ungava Bay
Labrador
South
North
Baffin
Nunavik
West
East
FMI IPI SAP KOA CHU KOV AVA LHO PRZ DUQ BDE FRM DOU PAY CHR HAB VOL FEU BER GEO AKI KAM ANA
K = 9
K = 10
K = 11
K = 12
K = 13
K = 14
K = 15

Figure S4: Loadings of the environmental factors on each axis of the PCA.

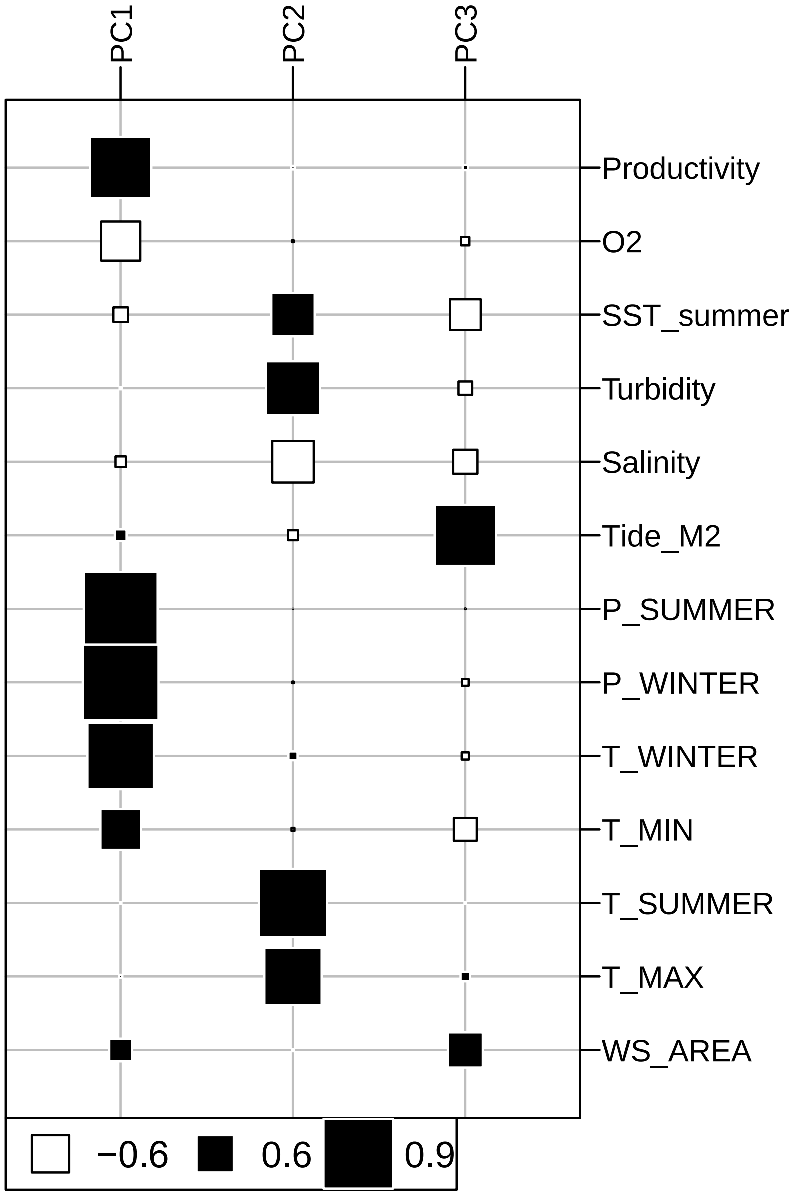

Figure S5: Sampling site scores for environmental components. The components correspond to the first 3 axis of a principal component analysis (PCA) of environmental variables. Scores are displayed in a color range of blue (lowest) to white (zero) to red (highest).

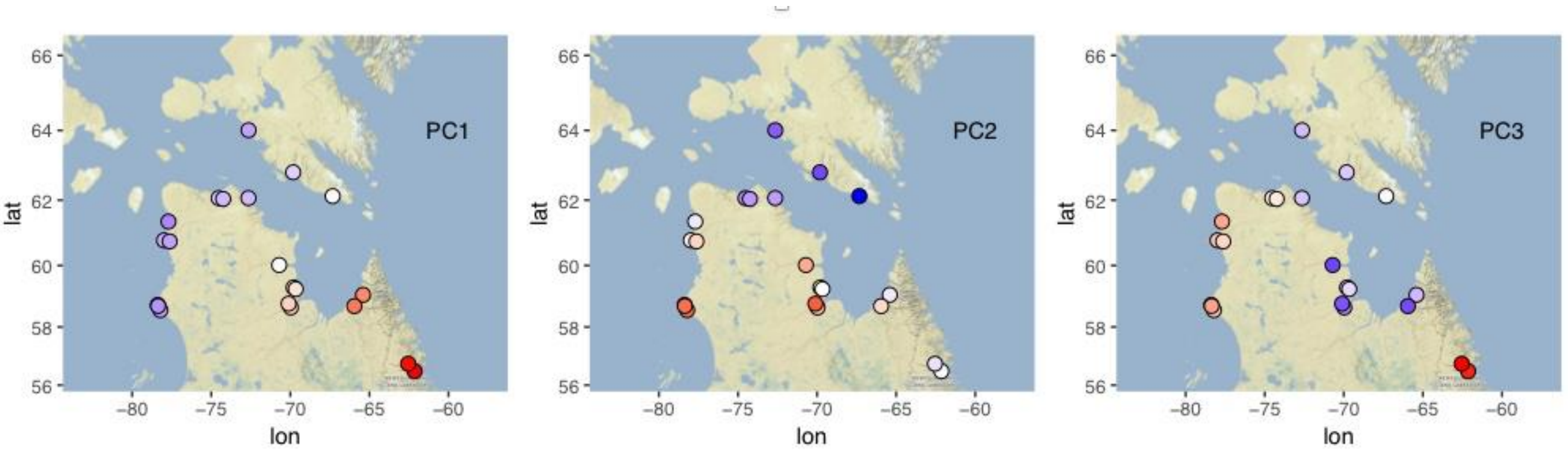

Figure S6: Values of distance-based Moran's eigenvectormap (dbMEM) components associated to positive spatial autocorrelation, displayed in a color range of yellow (lowest) to red (highest).

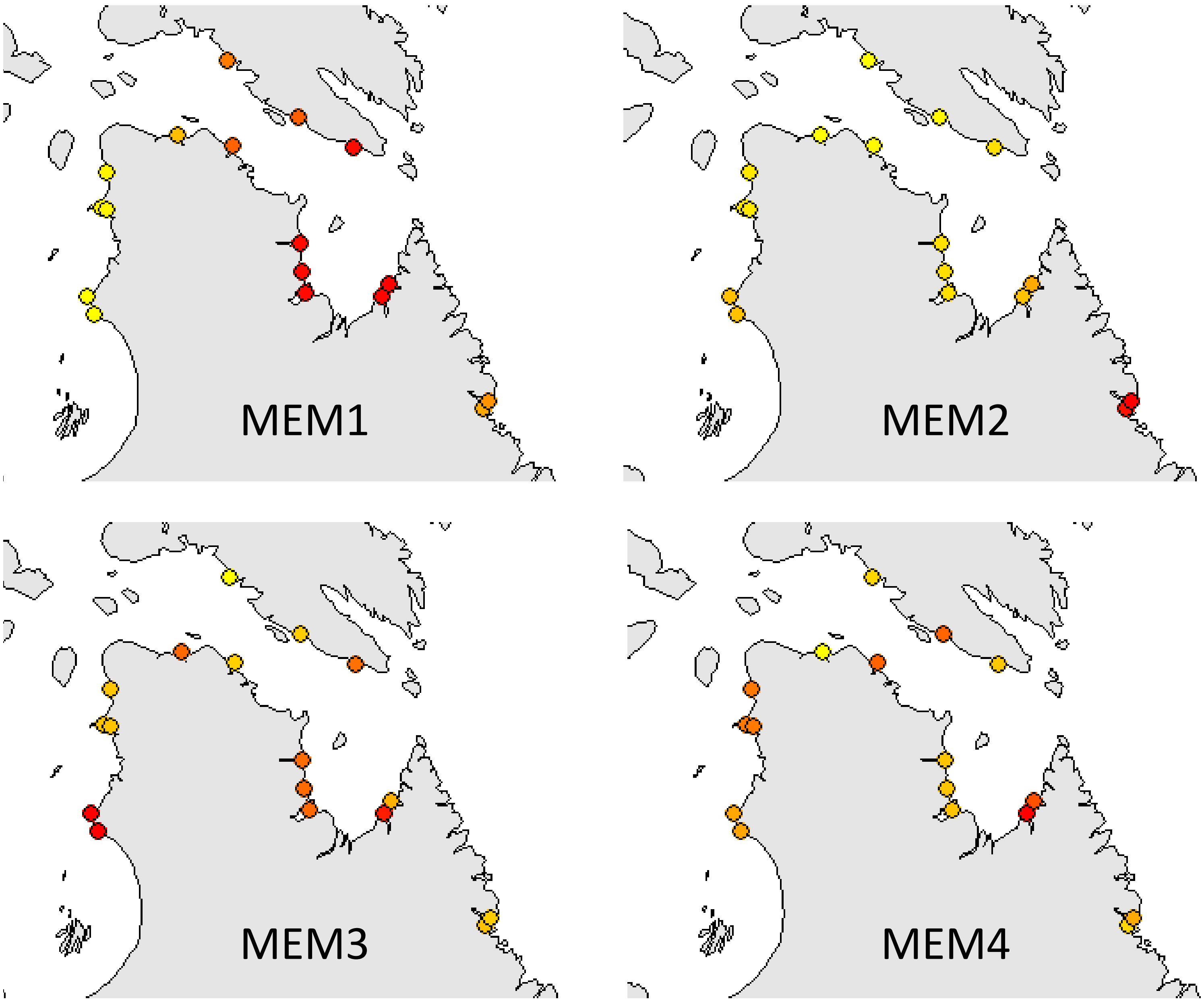

Figure S7: Absolute Z-score on the three first axis of a partial redundancy analysis (pRDA) based on 18,112 SNP markers. Dashed lines indicate significance thresholds (Z = 3.5, p = 0.0005). Candidate SNPs are colored according to the environmental component most correlated with allele frequencies.

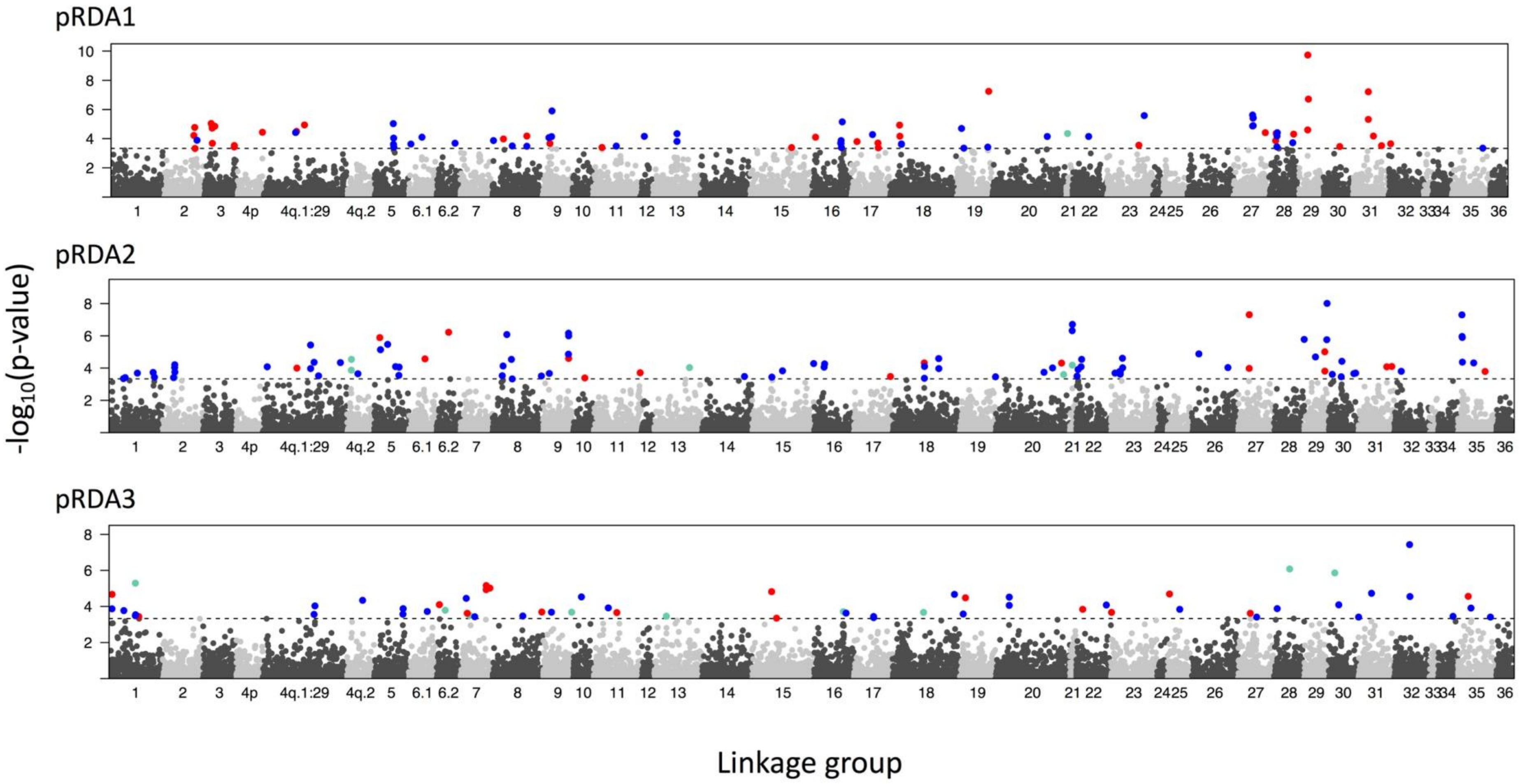

Figure S8: Triplots for (A) axes 1 and 2, and (B) axes 2 and 3 in a RDA excluding spatial components. The dark grey cloud of points at the center of each plot represents the SNPs, and coloured points represent sampling sites with color coding by region. Triplots are magnified to highlight SNP loadings on (C) RDA axes 1 and 2, and (D) axes 1 and 3. Candidate SNPs are shown as colored points with coding by most correlated environmental predictor (see text for description of predictors). Vectors represent environmental predictors, according to the scales on top and right axes.

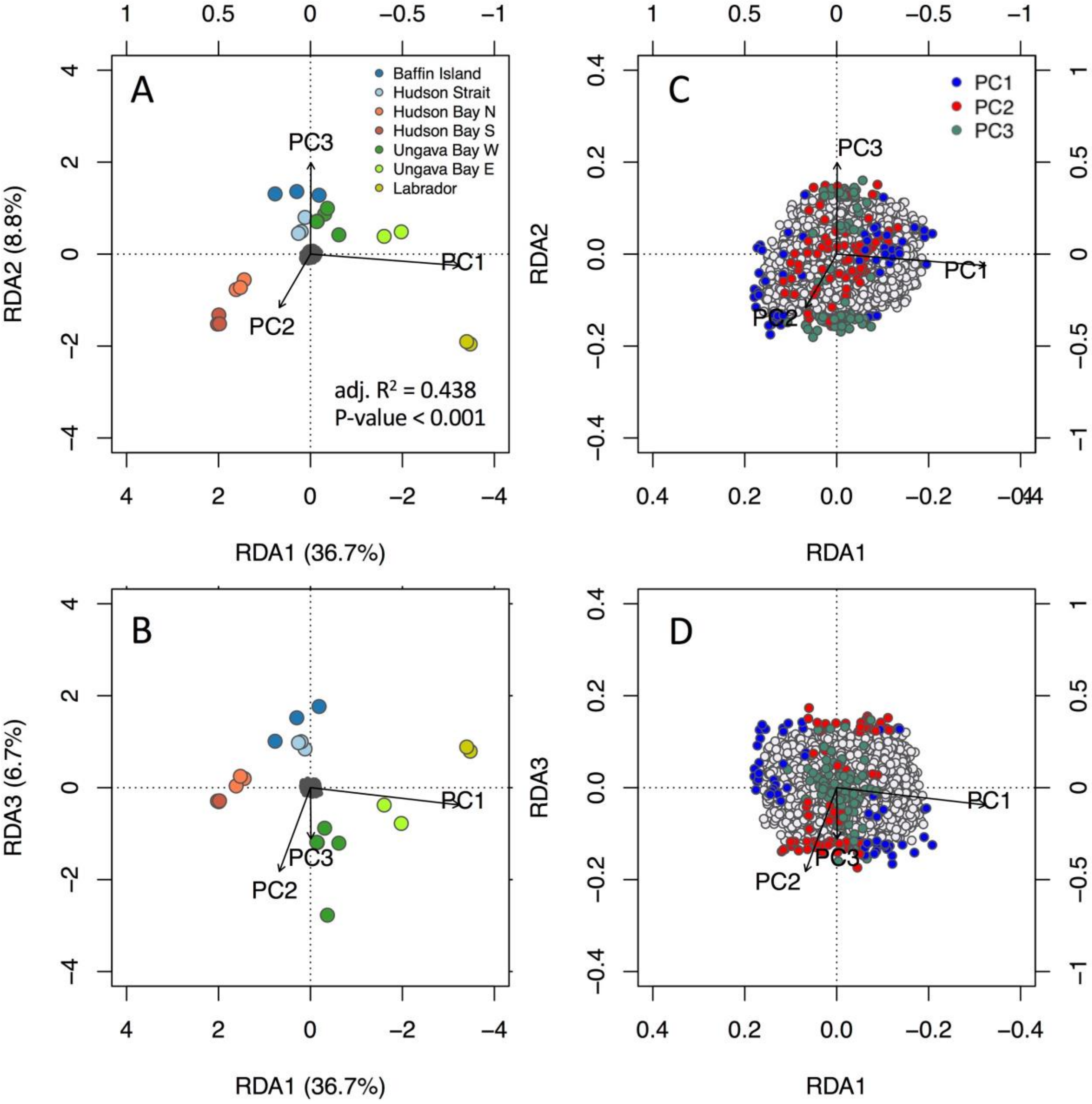

Figure S9: Absolute Z-score on the three first axis of a redundancy analysis (RDA) based on 18,112 SNP markers. Dashed lines indicate significance thresholds (Z = 3.5, p = 0.0005). Candidate SNPs are colored according to the environmental component most correlated with allele frequencies.

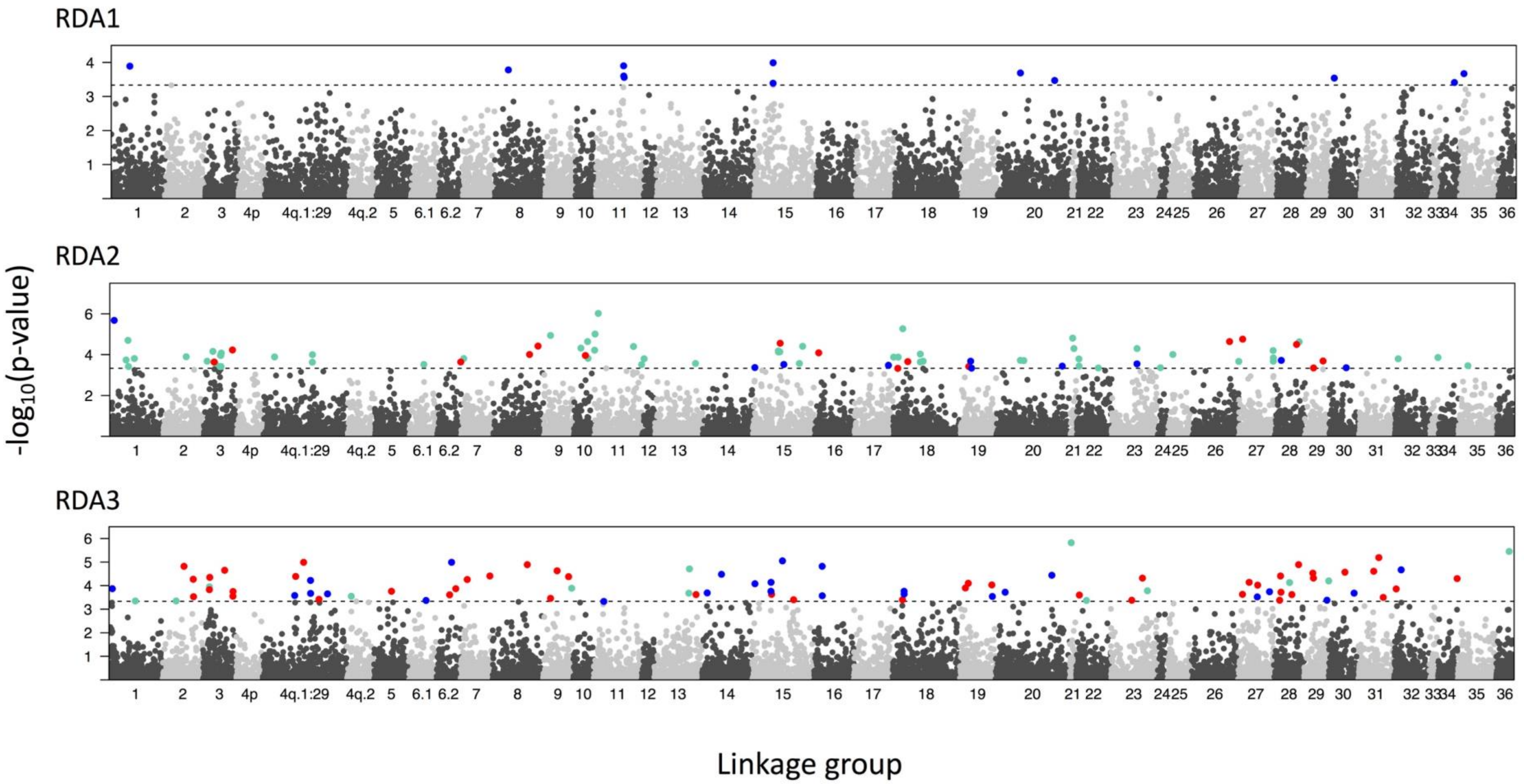

Figure S10: Q-values for 18,442 SNP markers and 3 environmental factors (PC1, PC2 and PC3) in a latent factor mixed model (LFMM). Dashed lines indicate significance thresholds (q-value < 0.01) and red dots mark candidate SNPs detected by at least one other GEA method.

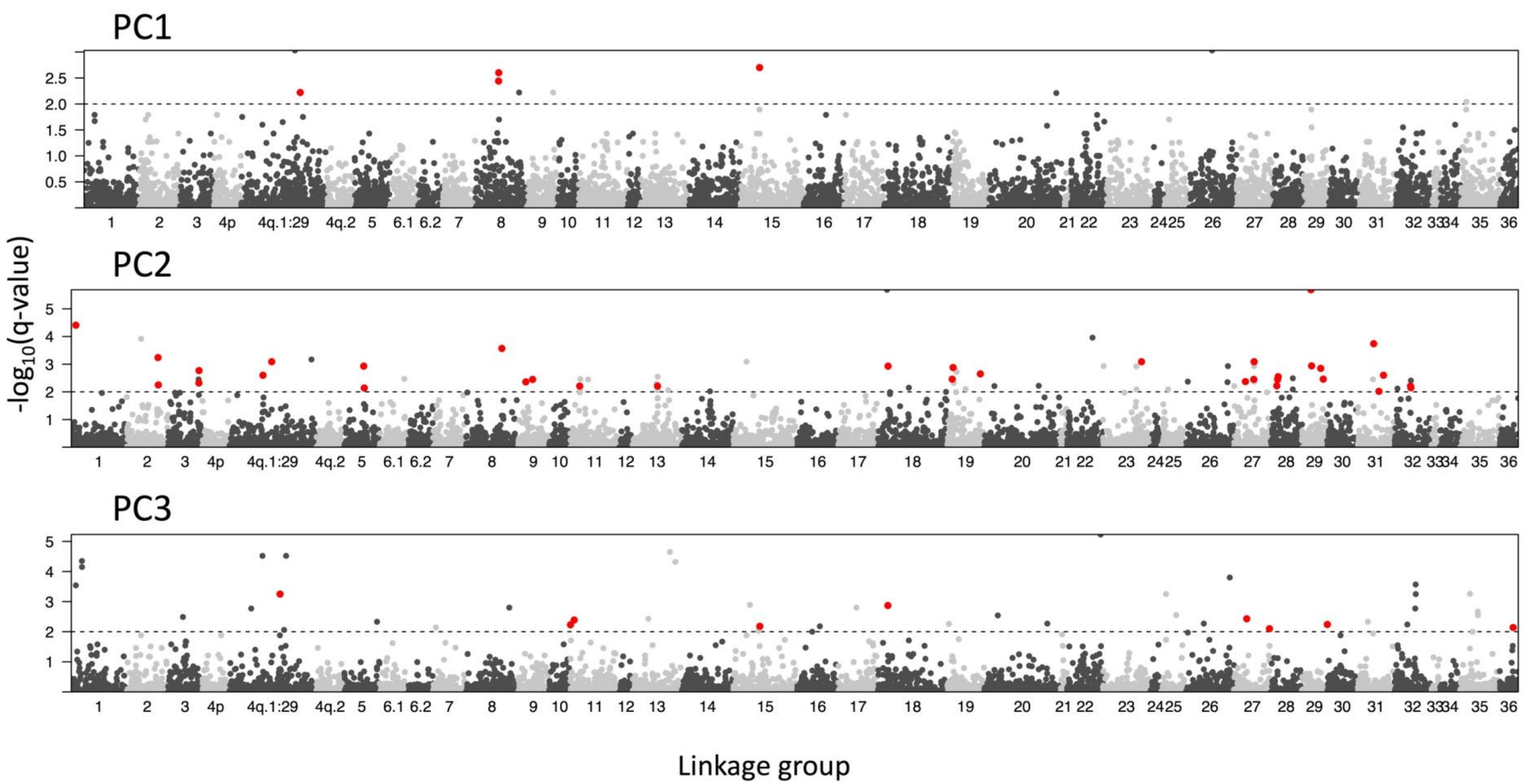

Figure S11: Median Bayesian Factor (BF) over 5 runs of the standard deviation model in Baypass for 18,442 SNP markers and 3 environmental factors (PC1, PC2 and PC3). Dashed lines indicate significance thresholds (BF > 10) and red dots mark candidate SNPs detected by at least one other GEA method.

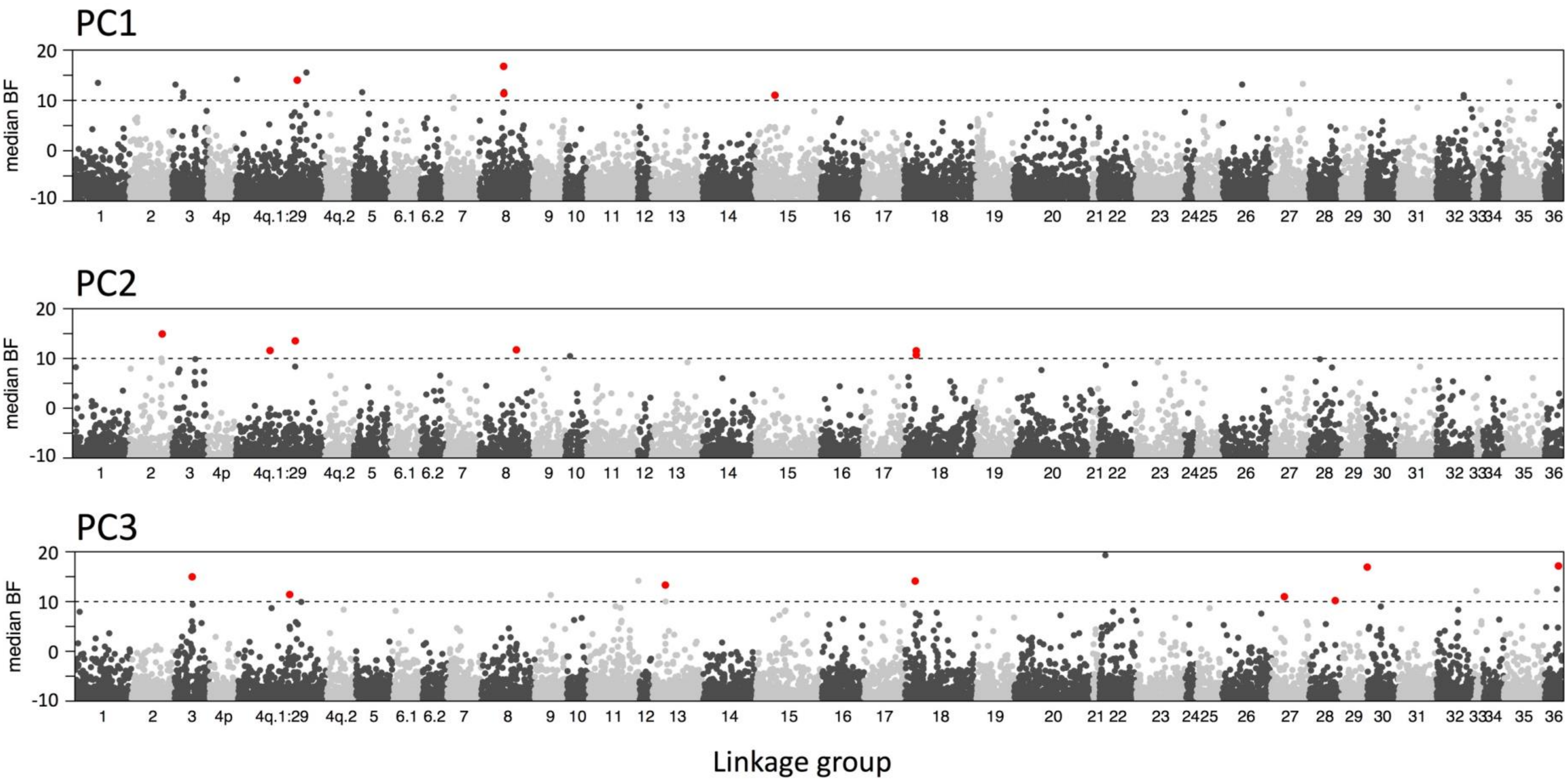

Figure S12: Frequencies of common (red) and rare (gray) alleles in sampling sites for candidate SNPs detected by at least two GEA methods. Environmental components associated to the SNP are listed for each method. Total: 75 SNP on 7 pages.

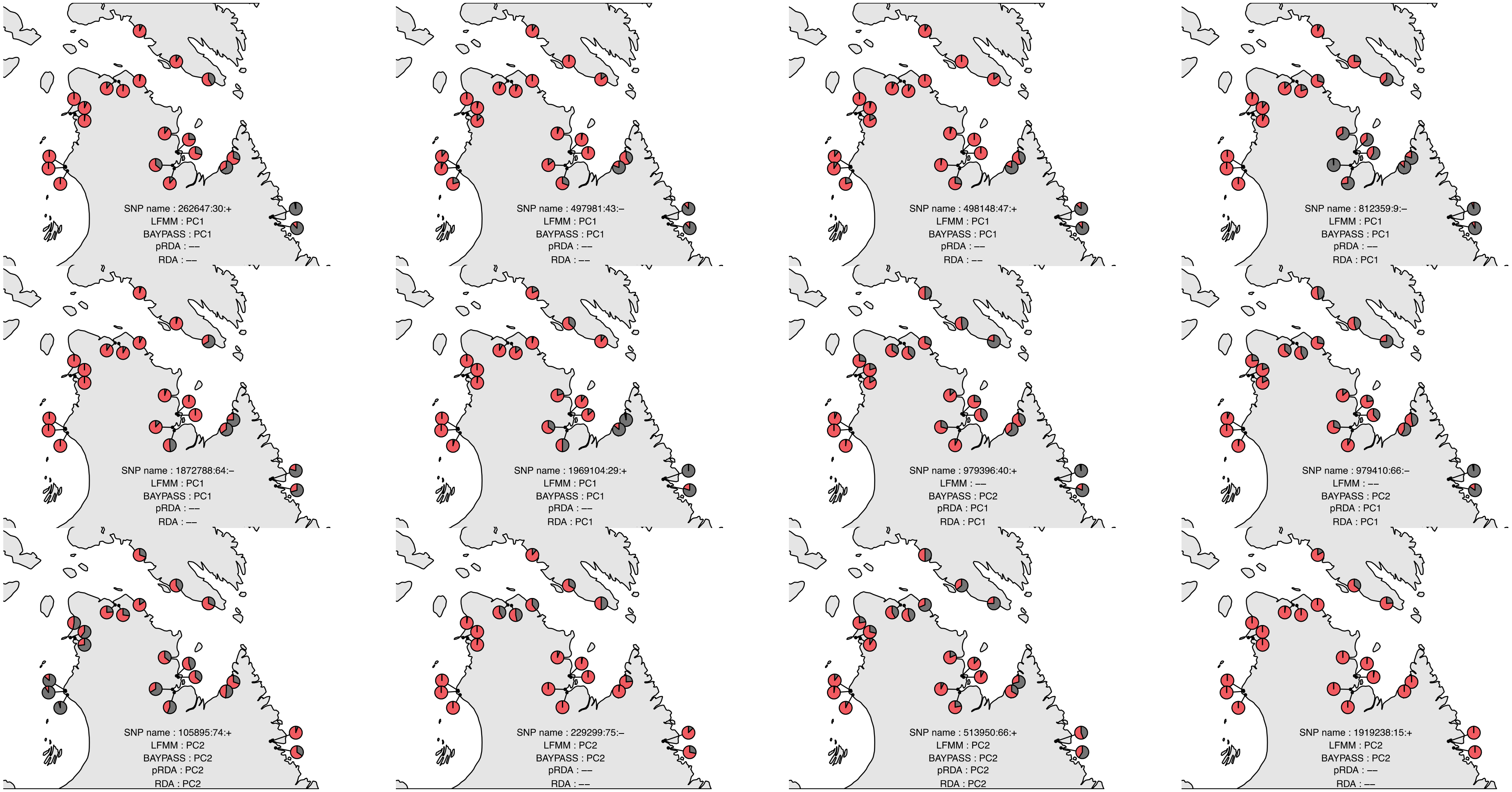

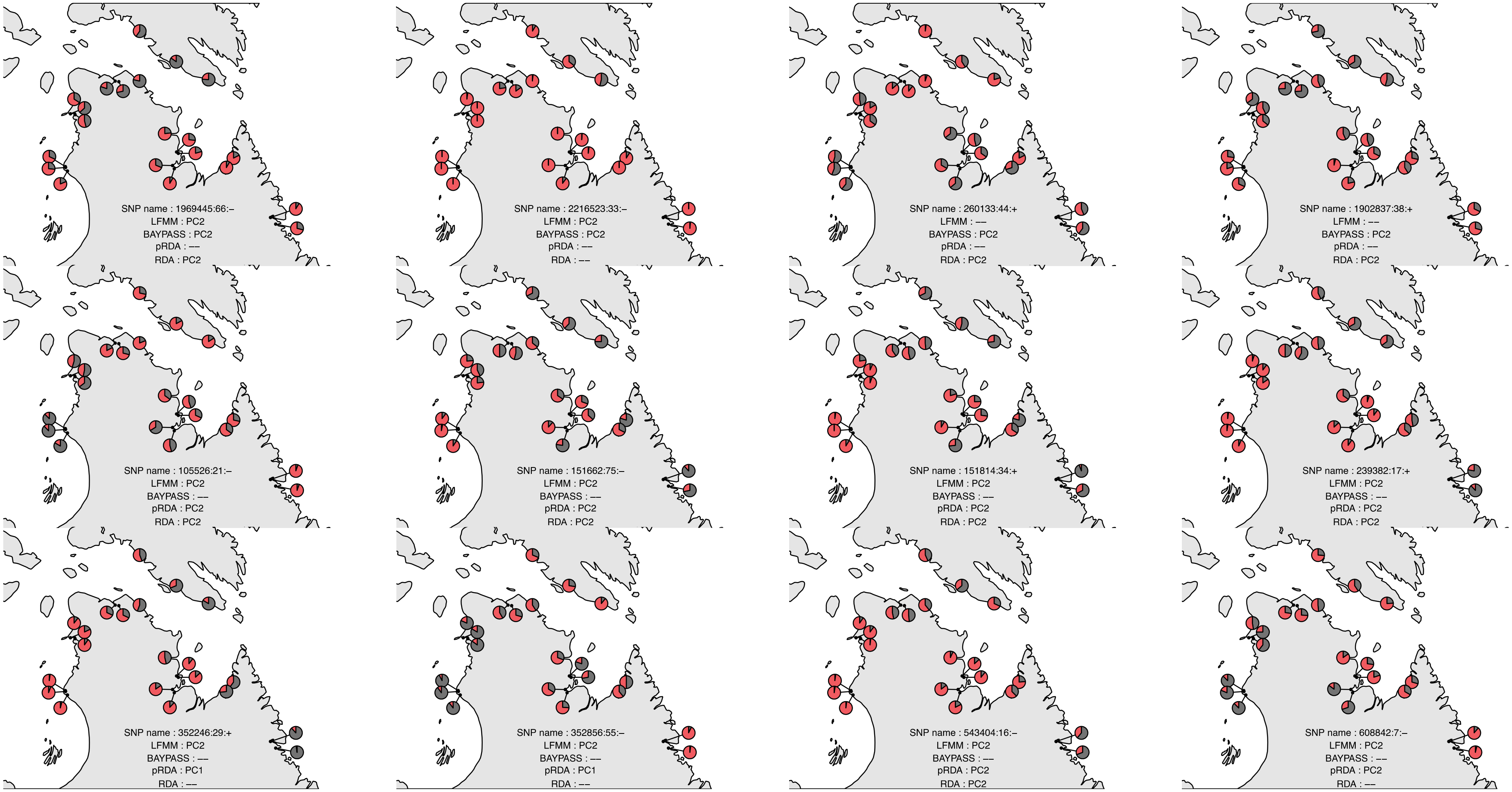

SNP name : 1969445:66:–
LFMM : PC2
BAYPASS : PC2
pRDA : ––
RDA : PC2
SNP name : 2216523:33:–
LFMM : PC2
BAYPASS : PC2
pRDA : ––
RDA : ––
SNP name : 260133:44:+
LFMM : ––
BAYPASS : PC2
pRDA : ––
RDA : PC2
SNP name : 1902837:38:+
LFMM : ––
BAYPASS : PC2
pRDA : ––
RDA : PC2
SNP name : 105526:21:–
LFMM : PC2
BAYPASS : ––
pRDA : PC2
RDA : PC2
SNP name : 151662:75:–
LFMM : PC2
BAYPASS : ––
pRDA : PC2
RDA : PC2
SNP name : 151814:34:+
LFMM : PC2
BAYPASS : ––
pRDA : PC2
RDA : PC2
SNP name : 239382:17:+
LFMM : PC2
BAYPASS : ––
pRDA : PC2
RDA : PC2
SNP name : 352246:29:+
LFMM : PC2
BAYPASS : ––
pRDA : PC1
RDA : ––
SNP name : 352856:55:–
LFMM : PC2
BAYPASS : ––
pRDA : PC1
RDA : ––
SNP name : 543404:16:–
LFMM : PC2
BAYPASS : ––
pRDA : PC2
RDA : PC2
SNP name : 608842:7:–
LFMM : PC2
BAYPASS : ––
pRDA : PC2
RDA : ––

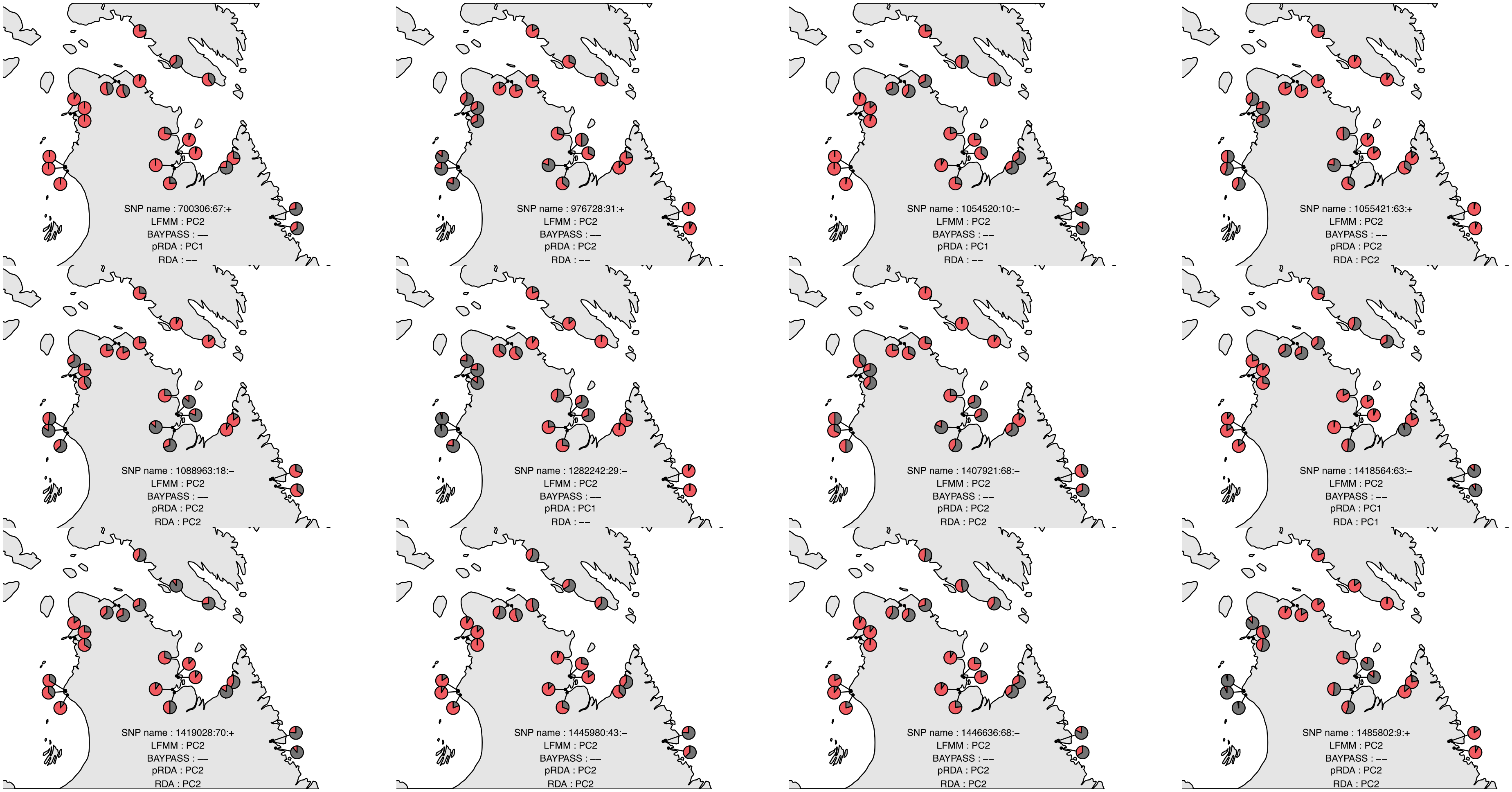
SNP name : 700306:67:+
LFMM : PC2
BAYPASS : ––
pRDA : PC1
RDA : ––
SNP name : 976728:31:+
LFMM : PC2
BAYPASS : ––
pRDA : PC2
RDA : ––
SNP name : 1054520:10:–
LFMM : PC2
BAYPASS : ––
pRDA : PC1
RDA : ––
SNP name : 1055421:63:+
LFMM : PC2
BAYPASS : ––
pRDA : PC2
RDA : PC2
SNP name : 1088963:18:–
LFMM : PC2
BAYPASS : ––
pRDA : PC2
RDA : PC2
SNP name : 1282242:29:–
LFMM : PC2
BAYPASS : ––
pRDA : PC1
RDA : ––
SNP name : 1407921:68:–
LFMM : PC2
BAYPASS : ––
pRDA : PC2
RDA : PC2
SNP name : 1418564:63:–
LFMM : PC2
BAYPASS : ––
pRDA : PC1
RDA : PC1
SNP name : 1419028:70:+
LFMM : PC2
BAYPASS : ––
pRDA : PC2
RDA : PC2
SNP name : 1445980:43:–
LFMM : PC2
BAYPASS : ––
pRDA : PC2
RDA : PC2
SNP name : 1446636:68:–
LFMM : PC2
BAYPASS : ––
pRDA : PC2
RDA : PC2
SNP name : 1485802:9:+
LFMM : PC2
BAYPASS : ––
pRDA : PC2
RDA : PC2

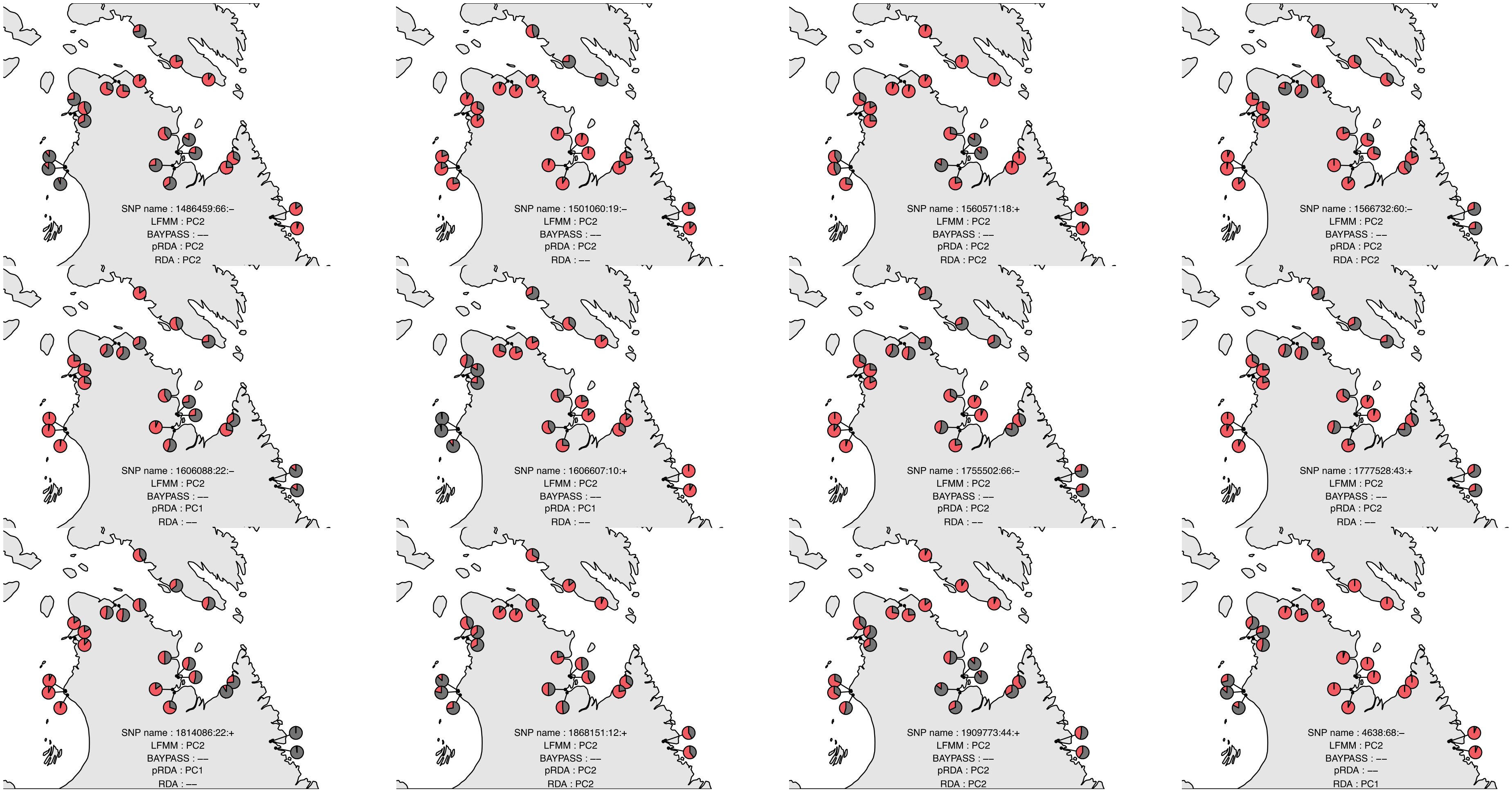
SNP name : 1486459:66:–
LFMM : PC2
BAYPASS : ––
pRDA : PC2
RDA : PC2
SNP name : 1501060:19:–
LFMM : PC2
BAYPASS : ––
pRDA : PC2
RDA : ––
SNP name : 1560571:18:+
LFMM : PC2
BAYPASS : ––
pRDA : PC2
RDA : PC2
SNP name : 1566732:60:–
LFMM : PC2
BAYPASS : ––
pRDA : PC2
RDA : PC2
SNP name : 1606088:22:–
LFMM : PC2
BAYPASS : ––
pRDA : PC1
RDA : ––
SNP name : 1606607:10:+
LFMM : PC2
BAYPASS : ––
pRDA : PC1
RDA : ––
SNP name : 1755502:66:–
LFMM : PC2
BAYPASS : ––
pRDA : PC2
RDA : ––
SNP name : 1777528:43:+
LFMM : PC2
BAYPASS : ––
pRDA : PC2
RDA : ––
SNP name : 1814086:22:+
LFMM : PC2
BAYPASS : ––
pRDA : PC1
RDA : ––
SNP name : 1868151:12:+
LFMM : PC2
BAYPASS : ––
pRDA : PC2
RDA : PC2
SNP name : 1909773:44:+
LFMM : PC2
BAYPASS : ––
pRDA : PC2
RDA : PC2
SNP name : 4638:68:–
LFMM : PC2
BAYPASS : ––
pRDA : ––
RDA : PC1

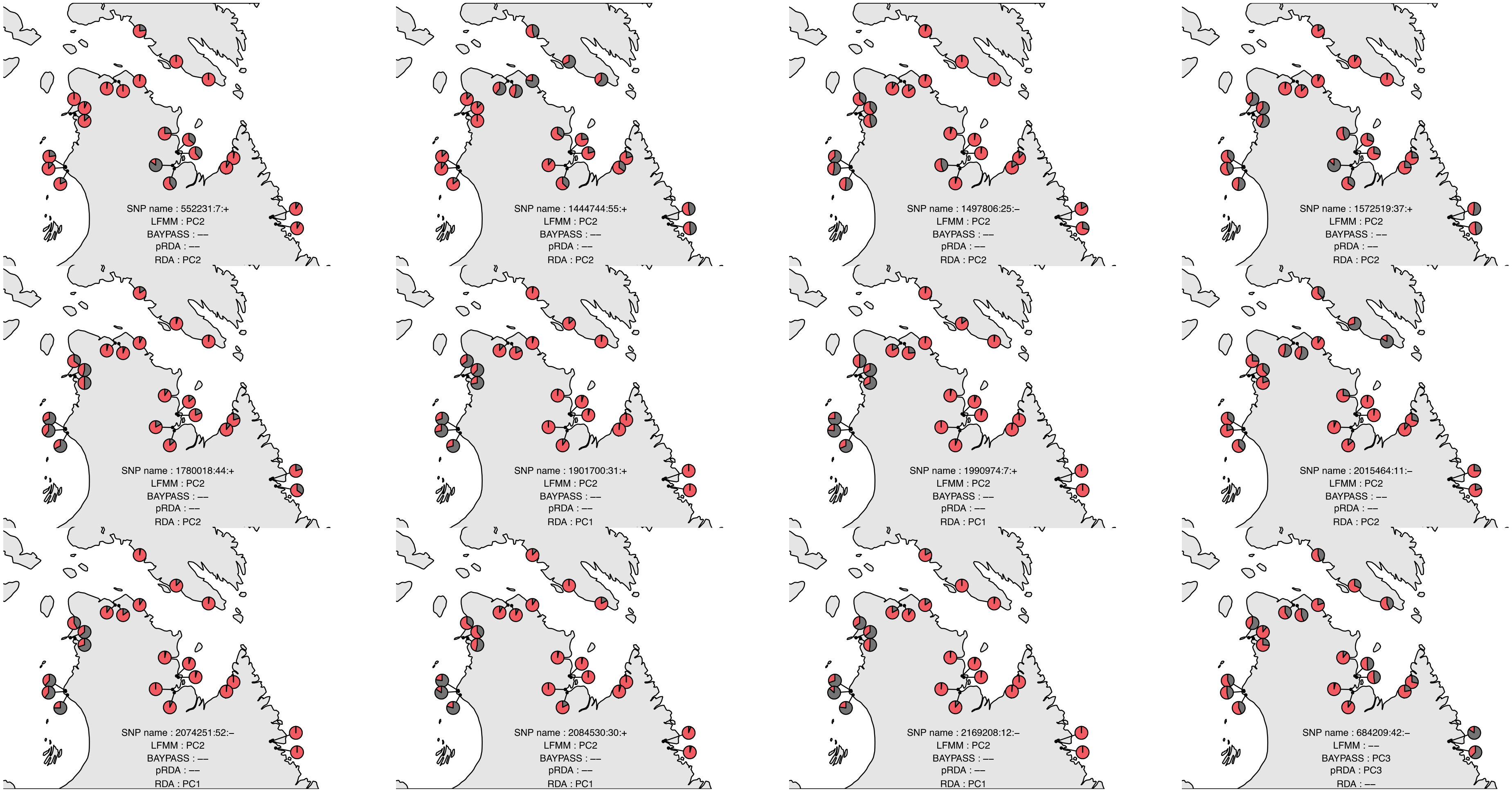
SNP name : 552231:7:+
LFMM : PC2
BAYPASS : ––
pRDA : ––
RDA : PC2
SNP name : 1444744:55:+
LFMM : PC2
BAYPASS : ––
pRDA : ––
RDA : PC2
SNP name : 1497806:25:–
LFMM : PC2
BAYPASS : ––
pRDA : ––
RDA : PC2
SNP name : 1572519:37:+
LFMM : PC2
BAYPASS : ––
pRDA : ––
RDA : PC2
SNP name : 1780018:44:+
LFMM : PC2
BAYPASS : ––
pRDA : ––
RDA : PC2
SNP name : 1901700:31:+
LFMM : PC2
BAYPASS : ––
pRDA : ––
RDA : PC1
SNP name : 1990974:7:+
LFMM : PC2
BAYPASS : ––
pRDA : ––
RDA : PC1
SNP name : 2015464:11:–
LFMM : PC2
BAYPASS : ––
pRDA : ––
RDA : PC2
SNP name : 2074251:52:–
LFMM : PC2
BAYPASS : ––
pRDA : ––
RDA : PC1
SNP name : 2084530:30:+
LFMM : PC2
BAYPASS : ––
pRDA : ––
RDA : PC1
SNP name : 2169208:12:–
LFMM : PC2
BAYPASS : ––
pRDA : ––
RDA : PC1
SNP name : 684209:42:–
LFMM : ––
BAYPASS : PC3
pRDA : PC3
RDA : ––

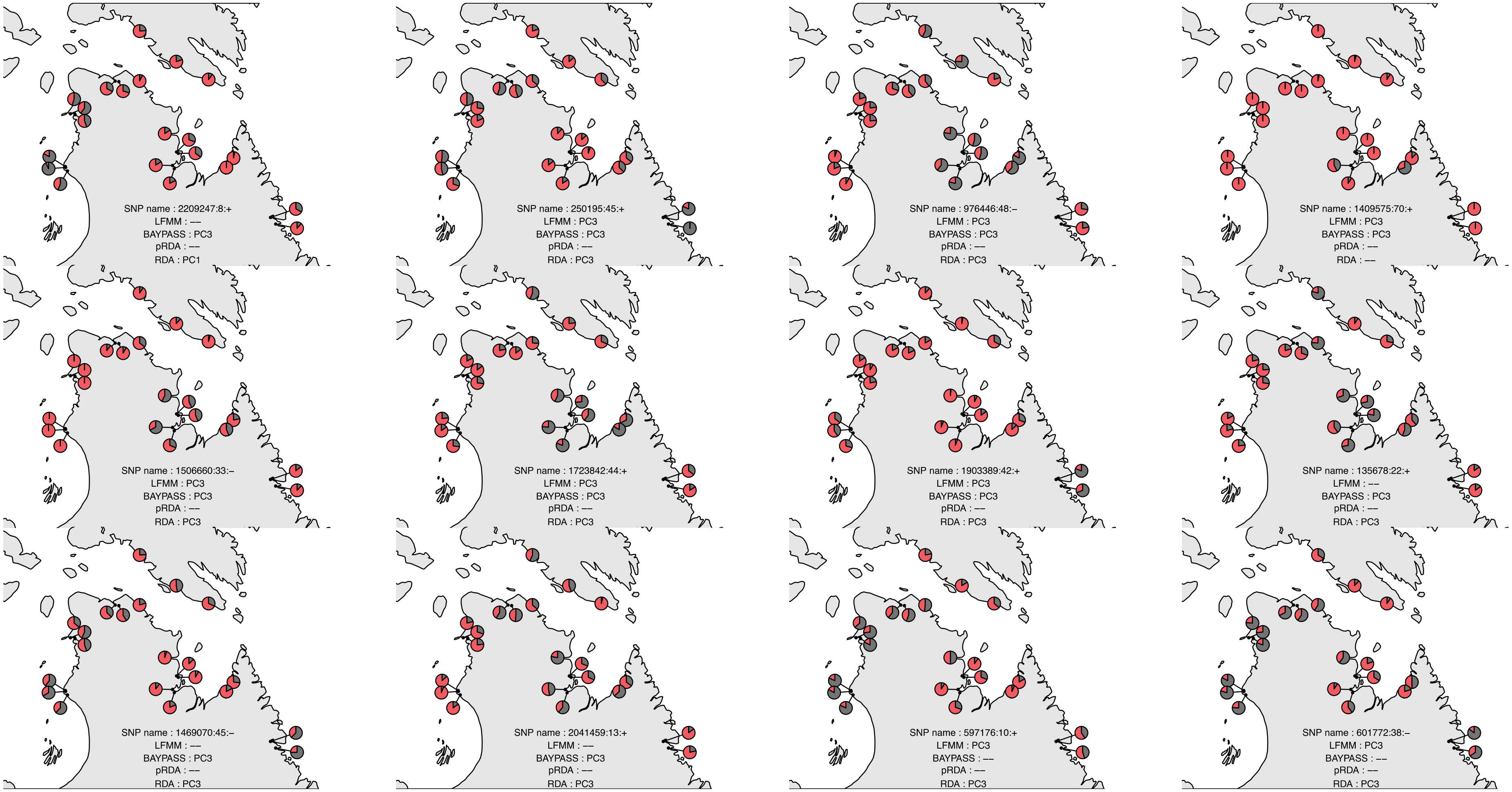
SNP name : 2209247:8:+
LFMM : ––
BAYPASS : PC3
pRDA : ––
RDA : PC1
SNP name : 250195:45:+
LFMM : PC3
BAYPASS : PC3
pRDA : ––
RDA : PC3
SNP name : 976446:48:–
LFMM : PC3
BAYPASS : PC3
pRDA : ––
RDA : PC3
SNP name : 1409575:70:+
LFMM : PC3
BAYPASS : PC3
pRDA : ––
RDA : ––
SNP name : 1506660:33:–
LFMM : PC3
BAYPASS : PC3
pRDA : ––
RDA : PC3
SNP name : 1723842:44:+
LFMM : PC3
BAYPASS : PC3
pRDA : ––
RDA : PC3
SNP name : 1903389:42:+
LFMM : PC3
BAYPASS : PC3
pRDA : ––
RDA : PC3
SNP name : 135678:22:+
LFMM : ––
BAYPASS : PC3
pRDA : ––
RDA : PC3
SNP name : 1469070:45:–
LFMM : ––
BAYPASS : PC3
pRDA : ––
RDA : PC3
SNP name : 2041459:13:+
LFMM : ––
BAYPASS : PC3
pRDA : ––
RDA : PC3
SNP name : 597176:10:+
LFMM : PC3
BAYPASS : ––
pRDA : ––
RDA : PC3
SNP name : 601772:38:–
LFMM : PC3
BAYPASS : ––
pRDA : ––
RDA : PC3

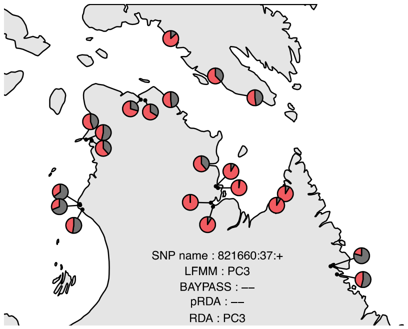
SNP name : 821660:37:+
LFMM : PC3
BAYPASS : ––
pRDA : ––
RDA : PC3

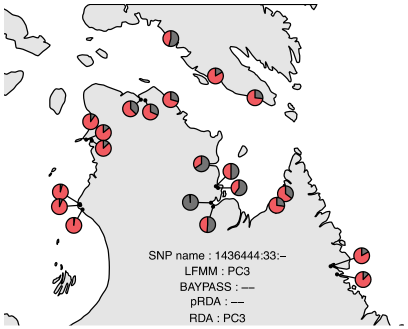
SNP name : 1436444:33:–
LFMM : PC3
BAYPASS : ––
pRDA : ––
RDA : PC3

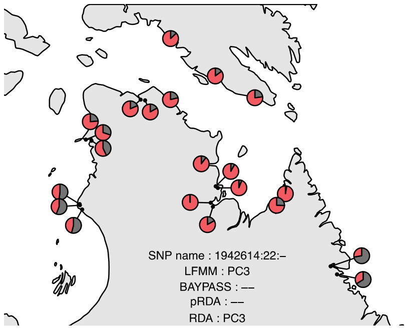
SNP name : 1942614:22:–
LFMM : PC3
BAYPASS : ––
pRDA : ––
RDA : PC3
